# Mesophasic organization of GABA_A_ receptors in hippocampal inhibitory synapse

**DOI:** 10.1101/2020.01.06.895425

**Authors:** Yun-Tao Liu, Chang-Lu Tao, Xiaokang Zhang, Lei Qi, Rong Sun, Pak-Ming Lau, Z. Hong Zhou, Guo-Qiang Bi

## Abstract

Information processing in the brain depends on synaptic transmission and plasticity, which in turn require specialized organization of neurotransmitter receptors and scaffolding proteins within the postsynaptic density (PSD). However, how these molecules are organized *in situ* remains largely elusive, limiting our mechanistic understanding of synaptic formation and functions. Here, we have developed template-free classification of over-sampled sub-tomograms to analyze cryo-electron tomograms of hippocampal synapses, enabling us to identify type-A γ-aminobutyric acid receptor (GABA_A_R) in inhibitory synapses and determine its *in situ* structure at 19 Å resolution. We found that these receptors are organized hierarchically: from GABA_A_R super-complexes with a fixed 11-nm inter-receptor distance but variable relative angles, through semi-ordered two-dimensional receptor networks with reduced Voronoi entropy, to mesophasic assembly with a sharp phase boundary. This assembly aligns with condensates of postsynaptic scaffolding proteins and putative presynaptic vesicle release sites. Such mesophasic self-organization may allow synapses to achieve a “Goldilocks” state with a delicate balance between stability and flexibility, enabling both reliability and plasticity in information processing.

## Introduction

Neuronal synapses are intricate communication devices, operating as fundamental building blocks underlying virtually all brain functions^1–4^. An essential part of the synapse is the lipid-bound, proteinaceous postsynaptic density (PSD), in which neurotransmitter receptors and other synaptic proteins are concentrated^5–8^. The specialized organization of the PSD is critical for the efficacy of synaptic transmission^9,10^. Meanwhile, the reorganization of receptors and other PSD proteins is widely known as a mechanism of synaptic plasticity, which in turn underlies many cognitive functions such as learning and memory^11,12^.

Different forms of PSD organization have been proposed, including meshwork based on electron microscopy and biochemical assays^13–15^, nano-domains based on super-resolution optical imaging^9,10,16–18^, and liquid condensate based on *in vitro* PSD mixing assay^19,20^. However, the PSD is heterogeneous and pleomorphic, and their protein components are small in size, presenting considerable challenges for resolving its molecular organization. For example, even super-resolution optical imaging can only describe synaptic organizations at the precision of protein clusters with its ∼20 nm resolution^16,17,21^. Electron microscopy, although with higher resolution, lacks molecular specificity, thus hindering the ability to identify synaptic receptors and other proteins inside synapses. These synaptic molecules, such as type-A γ-aminobutyric acid receptors (GABA_A_Rs), are often small and surrounded by the crowded cellular environment. Consequently, how individual PSD molecules are organized *in situ* is largely unknown, limiting our understanding of molecular mechanisms underlying synaptic formation and functions.

Here, we employed the state-of-the-art cryo electron tomography (cryoET) with Volta phase plate and direct electron detector to obtain structures of neuronal synapses in their native conditions. In order to automatically identify neurotransmitter receptors inside synapses without the need of labeling, we developed a method of template-free classification with uniformly oversampled sub-tomograms on the membrane. With this method, we obtained an *in situ* structure of GABA_A_R at 19 Å resolution and discovered a hierarchical organization of GABA_A_Rs within the PSD, establishing the structural basis for synaptic transmission and plasticity.

## Results

### Identification of GABA_A_Rs by oversampling and template-free classification

To understand the molecular organization of GABA_A_Rs *in situ*, we imaged synapses of cultured hippocampal neurons using cryoET with Volta phase plate (Supplementary Video 1). Taking advantage of correlative microscopy, we have shown that a thin sheet-like density parallel to the postsynaptic membrane is a defining feature of GABAergic inhibitory synapses^7^ (Fig. 1a). Following this criterion, we identified that 72 synapses in the tomograms we obtained are inhibitory synapses. Many particles visualized on the postsynaptic membrane in these synapses have shapes characteristic of pentameric GABA_A_R^22^ (Figs. 1b-e; Supplementary Video 2), which is the most abundant membrane protein species in GABAergic synapses^23,24^ (Extended Data Table 1). We thus assigned these particles as GABA_A_Rs on the native postsynaptic membrane. To automate the unbiased identification of GABA_A_Rs, we devised a systematic approach that uses oversampling of sub-tomograms to ensure inclusion of all particles existing on the postsynaptic membranes, and then classifies the oversampled sub-tomograms with template-free, Bayesian 3D classification method as implemented in Relion^25^ to sort out GABA_A_R particles from all the particles (Extended Data Figs. 1-4; see also Methods). The structure of GABA_A_R emerged during the iterative classification (Extended Data Fig. 1c). After eliminating duplicates, we sorted out 9,618 GABA_A_Rs from all 72 synapses (Extended Data Fig. 1b) and placed them back on the postsynaptic membranes to visualize their spatial distribution (Fig. 1d). After 3D refinement, a sub-tomogram average of *in situ* GABA_A_R was obtained at 19 Å resolution (Fig. 1f-g).

**Fig. 1.**
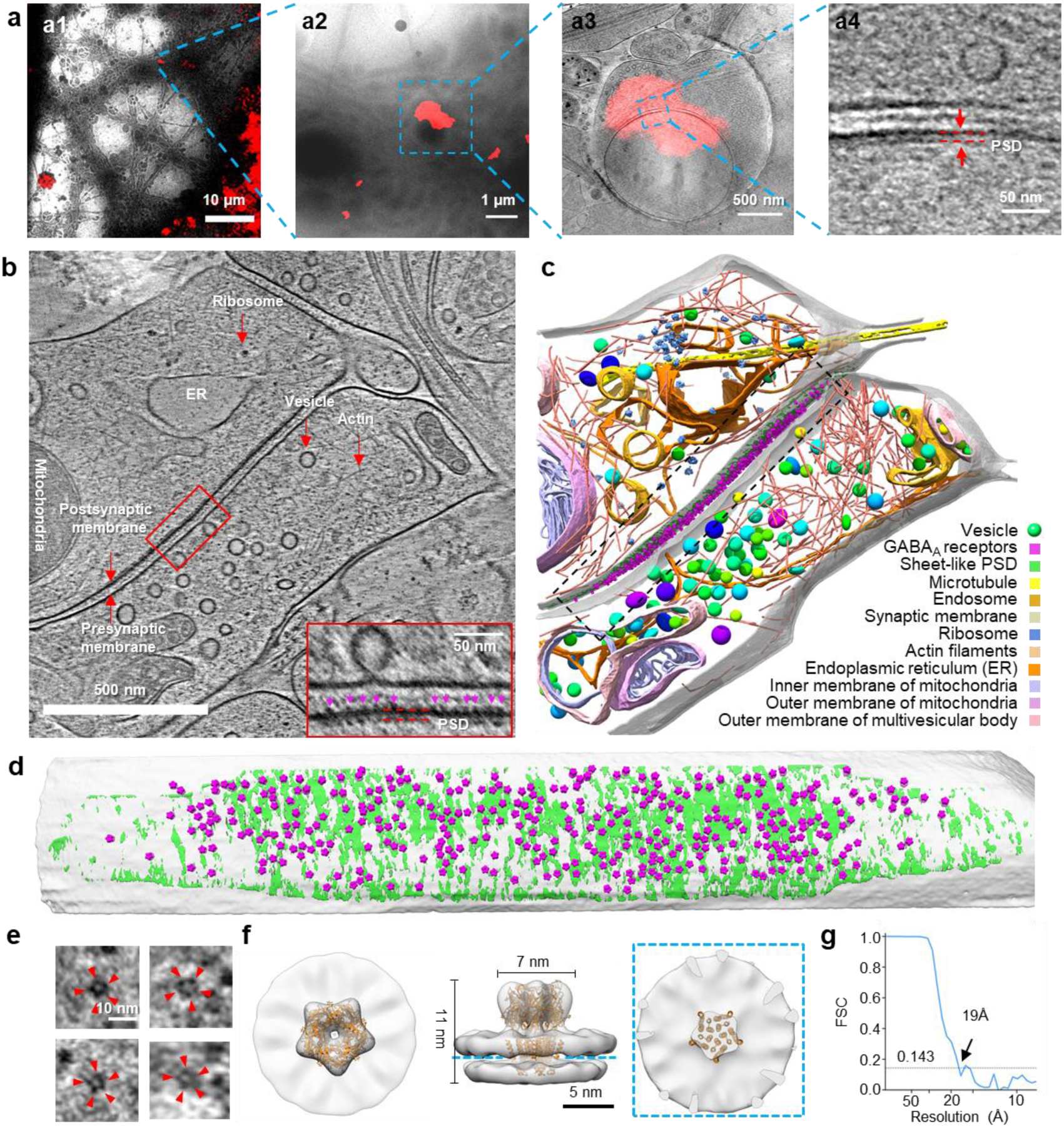
Identification and *in situ* structure of GABA_A_R in inhibitory synapse. **a,** Identification of inhibitory synapses with cryo-correlative light and electron microscopy. **a1,** Low magnification EM image superposed with fluorescence image of gephyrin-mCherry. **a2,** Zoomed-in view of (**a1**). **a3,** Electron tomographic slice superposed with fluorescence puncta. **a4,** Zoomed-in view of (**a3**) showing thin sheet-like PSD. **b,** A tomographic slice of an inhibitory synapse. Various subcellular components are labelled on the image. Inset: Zoomed-in view showing receptor densities (magenta arrowheads). **c,** 3D rendering of the tomogram shown in (**b**). **d,** Front view of GABA_A_Rs (purple), density of the scaffolding protein layer (green) on the postsynaptic membrane (transparent gray). **e,** Example tomographic slices of individual GABA_A_R in top view. Red arrowheads showing 5 blobs of GABA_A_R density. **f,** Sub-tomogram average of GABA_A_R fitted with crystal structure (orange ribbons)^22^. **g,** Fourier shell correlation of the GABA_A_R sub-tomogram average.

### *In situ* structure of GABA_A_R

The sub-tomogram average of GABA_A_R was ∼11 nm in length and ∼7 nm in width and had a central pore (Fig. 1f). Overall, our *in situ* structure matched the previously characterized structure of reconstituted GABA_A_R^22^, except that extra densities were found at the edges of the extracellular domain (Fig. 1f). These extra densities might represent additional glycans only existing in the native proteins expressed in neurons^26^. Densities for the membrane bilayer were also well resolved (Fig. 1f). The rough shape of the density for the transmembrane helices matched the atomic models of the reconstituted GABA_A_Rs^22,26–29^, with some slight difference that could be due to averaging of different subunits (Fig. 1f; Extended Data Fig. 5a). The intracellular loops (∼500 a.a, for 5 subunits, missing in atomic structures) were not observed in our reconstruction even at low threshold (Extended Data Fig. 5b), suggesting that those loops are intrinsically flexible even though they are likely to bind to postsynaptic scaffolding proteins *in situ*.

### Super-complex of GABA_A_Rs

With the GABA_A_Rs identified *in situ*, we next investigated their spatial organization on the postsynaptic membrane. By measuring the distance of each receptor to its neighbors, we found that the distributions of the first and the second nearest neighbor distances both peaked sharply at ∼11 nm (Fig. 2a-b), indicating that GABA_A_Rs tend to maintain a fixed distance with their neighboring receptors. Receptor concentration measured as number of particles/µm^2^ within the concentric rings around GABA_A_Rs also peaked at ∼11 nm (Fig. 2c), further supporting that 11 nm is a characteristic inter-receptor distance (IRD). At this distance, the concentration of GABA_A_R reached ∼4,000 µm^-2^, which was about twice of the plateau level that occurs just 5 nm away (Fig. 2c). This characteristic 11-nm IRD was consistently found in most (64 out of 72) synapses (Fig. 2d). The rest had generally fewer receptors and larger median IRDs (Fig. 2d), probably due to their immaturity in early synapse development. By selecting receptors and their neighboring receptors with 11±4 nm IRDs (Fig. 2e). we obtained a sub-tomogram average of GABA_A_R super-complex consisting of a pair of receptors (Fig. 2f). Moreover, classification of oversampled sub-tomograms without symmetry also yielded a class with a pair of receptor-like particles with ∼11 nm IRD (Extended Data Fig. 4). Thus, this IRD imposes a stringent constraint on the organization of GABA_A_Rs on the inhibitory postsynaptic membrane. In the averaged receptor pair super-complex, the pseudo 5-fold symmetry in both receptors was lost, suggesting that the relative rotation of each receptor was less constrained (Fig. 2f). Indeed, the distribution of the in-plane rotation angle (denoted as angle ω) of a receptor relative to the receptor pair axis was quite uniform (Fig. 2g). Reconstructing the receptor pairs with specific ω angles clearly restored the pseudo-5-fold symmetry of the corresponding receptor (Fig. 2h).

**Fig. 2.**
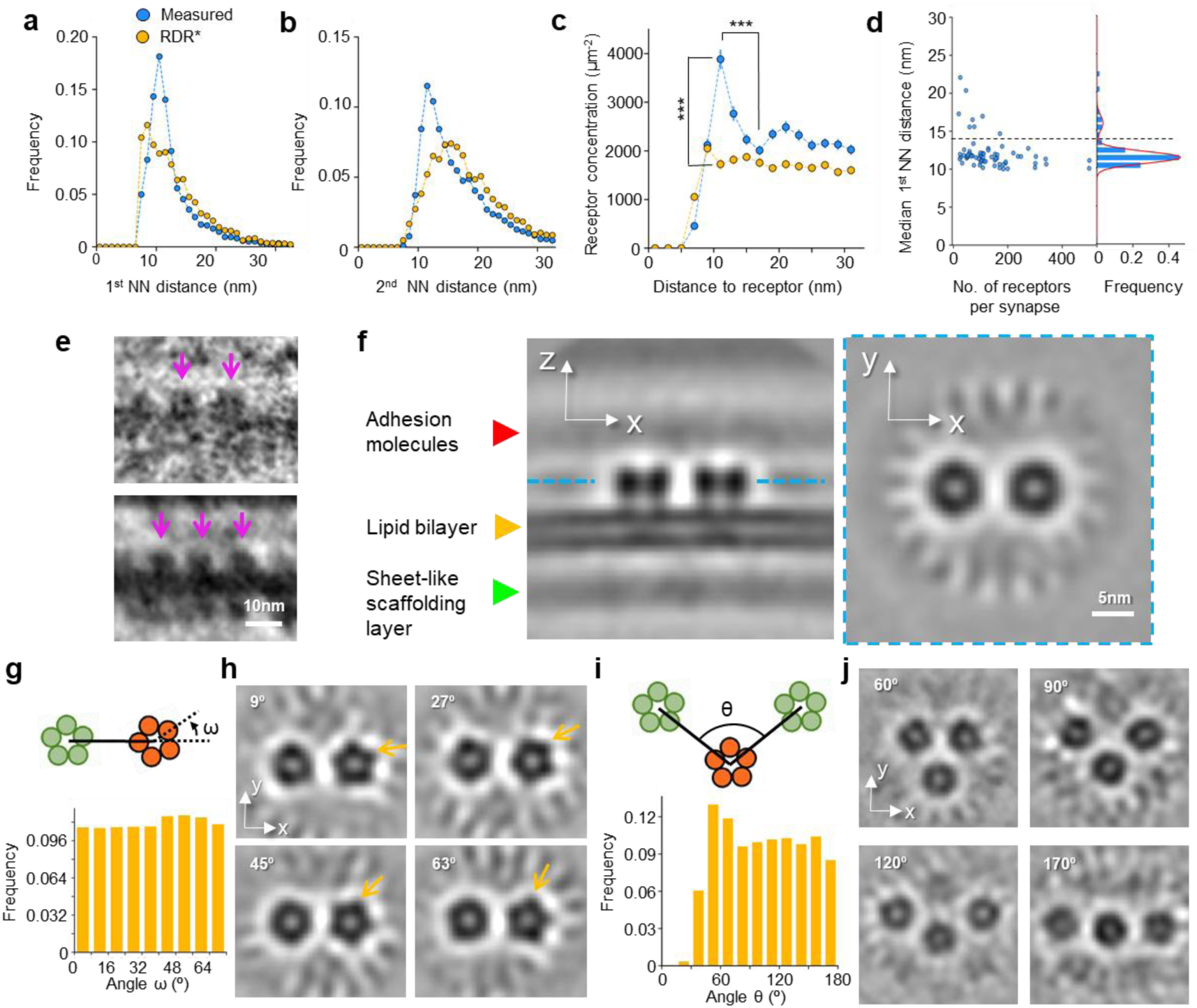
GABA_A_R super-complexes. **a, b,** The first (**a**) and second (**b**) nearest neighbor (NN) distances distribution of measured receptors (Measured) and randomly distributed receptors without overlap (RDR*, *i.e.* distance between any two receptors is larger than 7 nm). P<0.001, K-S test in both (**a**) and (**b**). **c,** Receptor concentration as a function of distance to a GABA_A_R (n=9,618). ***, P<0.001, two-tailed t-test. **d,** Left: Scatter plot of the number of receptors vs. the median first NN distance of each synapse. Right: Frequency distribution of median first NN distance, fitted with two Gaussian distributions (red curve). Dashed line shows the lowest point between two peaks. **e,** Examples of receptor pairs and triplets from original tomograms. Arrows point to receptors. **f,** Orthogonal slice views of the sub-tomogram average of receptor pairs. **g, i,** The distribution of relative rotation angles ω (**g**) and θ (**i**), as defined in respective diagrams. **h, j,** Sub-tomogram averages of receptor pairs with different ω (**h**) and receptor triplets with different θ (**j**). Error bars are SEM for all figures.

One GABA_A_R could also pair with two other receptors, forming a receptor triplet (Figs. 2e, i). In the triplet structure, whereas the distances between the neighboring receptors were constrained to ∼11 nm, the angle (denoted as angle θ) between the two arms of the triplet was unrestricted, with a rather uniform distribution ranging from 60° to 180° (Fig. 2i). The structures of receptor triplets with different θ angles could also be reconstructed (Fig. 2j). Thus, the near-neighbor organization of GABA_A_Rs is morphologically flexible with variable ω or θ angles, but topologically invariable with fixed IRD. This unique feature is characteristic of a mesophasic state, which is neither liquid that does not maintain inter-molecule distance, nor crystalline that has constant crystal angles.

### Two-dimensional networks of GABA_A_Rs

In addition to pairs and triplets of “linked” receptors, many receptors (26.1%) in fact had more than two 11-nm neighbors (Fig. 3a), and further organized into two-dimensional networks of various sizes and shapes (Fig. 3b). In the meantime, 20.0% receptors did not integrate into the network, hereafter defined as solitary receptors (Figs. 3a-b). The proportion of solitary receptors and mean size of the networks were independent of postsynaptic area or number of receptors in a synapse (Fig. 3c; Extended Data Fig. 6a), consistent with the idea that the function of these synapses could be altered independently either by changing number of receptors or by modifying the organization of postsynaptic protein network^30^.

**Fig. 3.**
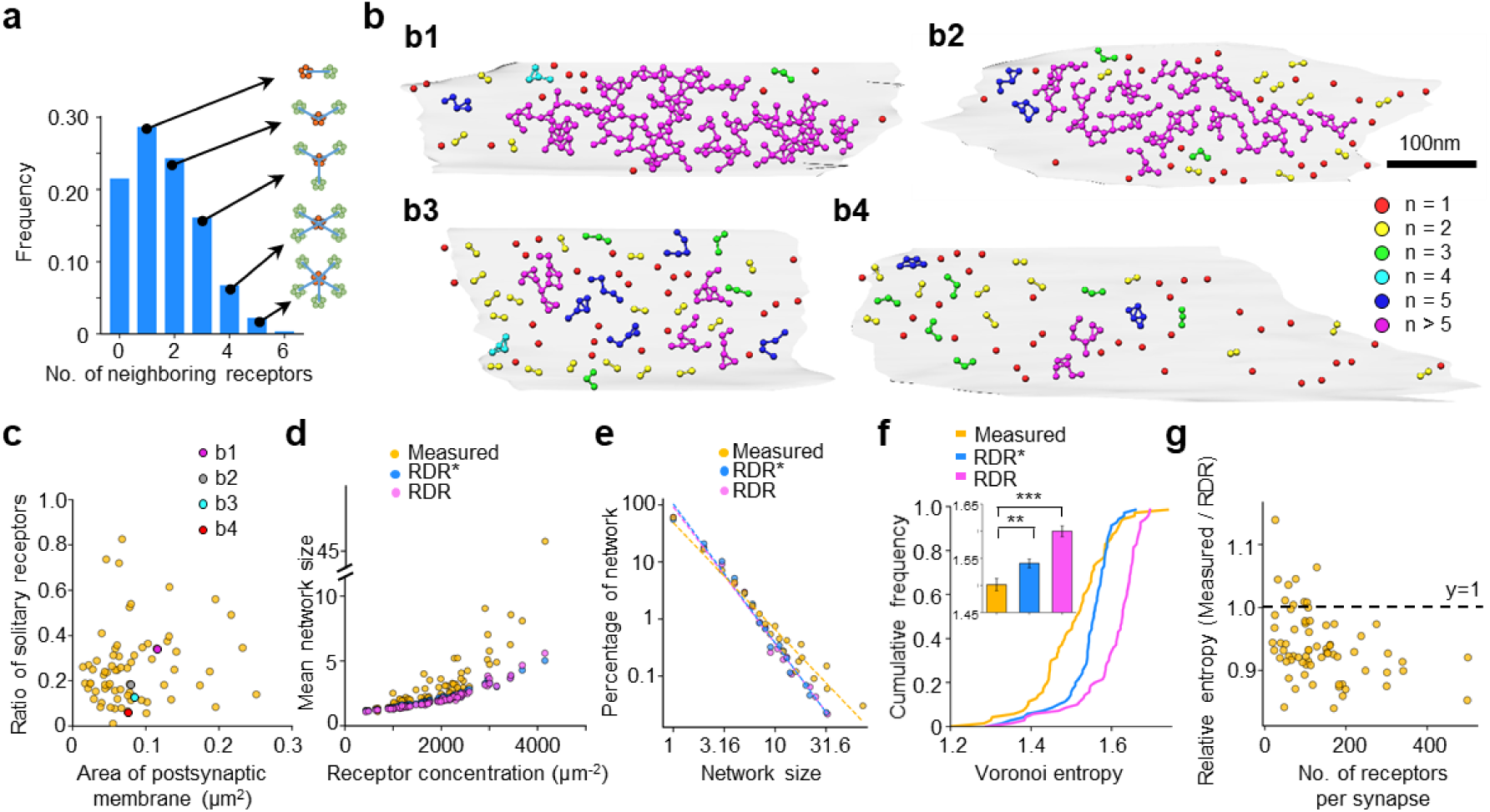
Two dimensional networks of GABA_A_Rs. **a,** Distribution of the number receptors with different numbers of 11-nm neighbors. **b,** Four examples (**b1-4**) of receptor network organization on the postsynaptic membrane. Color indicates network size (n, the number of receptor in a network). **c,** Scattered plot of the ratio of solitary receptors vs. the area of postsynaptic membrane for each synapse. Colored dots (magenta, gray, cyan and red) correspond to synapses in (**b**). **d,** Scattered plot of mean network size vs. receptor concentration for measured receptors (Measured), randomly distributed receptors (RDR), and randomly distributed receptors without overlap (RDR*). **e,** Power law distribution of network size in log-log plot. **f,** Cumulative frequency of Voronoi entropy for each synapse (n=70). Inset: Mean value of Voronoi entropy. **, P<0.01, ***, P<0.001, K-S test. **g,** Scattered plot of relative entropy (defined as Measured/RDR) vs the number of receptors for each synapse.

Intriguingly, the mean size of receptor networks in a synapse, when plotted against receptor concentration, was always larger than that for simulated randomly distributed receptors (RDR) or randomly distributed receptors without overlap (RDR*) (Fig. 3d). Furthermore, the overall distribution of network size followed power law (Fig. 3e; Extended Data Fig. 6b). The power-law exponent (1.87), representing the fractural dimension of receptor networks, was smaller than that for RDR (2.44) and RDR* (2.40) (Fig. 3e). These results suggest that receptor networks tend to “attract” more receptors to grow into larger networks, a property typically found in self-organizing processes near critical states^31^.

To quantify the degree of orderliness for the receptor organization, we calculated Voronoi entropy that measures information content in the Voronoi tessellation of the receptor localizations^32^ (Extended Data Fig. 6c). The Voronoi entropy becomes zero for a perfectly ordered structure, while for a fully random 2D distribution of points the value has been reported to be 1.71^33^. The Voronoi entropy for our measured receptor distribution was 1.50, smaller than that for RDR (1.60) and RDR* (1.55) (Fig. 3f). The smaller entropy for the measured receptors is likely to arise from the semi-ordered 2D networks. This Voronoi entropy value in between the entropy of crystal and liquid further suggests that the receptors organize in mesophasic state. This mesophasic state is apparently much more disorder than the liquid-crystalline state of acetylcholine receptors in neuromuscular junction^34,35^. This could potentially allow for rapid change in receptor organization to serve as a plasticity mechanism in GABAergic synapses. Several synapses (12.9%) had Voronoi entropy larger than that of RDR (Fig. 3g). They are mostly synapses with fewer receptors that were unable to establish semi-ordered organization.

### Mesophasic assembly of inhibitory PSD

The semi-ordered receptor networks presumably reflect a mesophasic state of the self-organized PSD. If this is the case, one would expect that the mesophasic PSD may separate from its aqueous environment with a phase boundary. To test this, a smoothed convex hull of all linked receptors (Fig. 4a1; Extended Data Figs. 7a-b) was constructed. Within this hull that enclosed about 66% of the postsynaptic membrane area (Extended Data Fig. 7c), the receptor concentration was high (∼3,000 µm^-2^) and relatively uniform. This concentration dropped steeply within ∼18 nm across the hull (Fig. 4b). Thus, the smoothed convex hull can indeed be considered as the phase-separating boundary of the mesophasic receptor assembly. Interestingly, the sharp boundary was characteristic only for the linked receptor, whereas the concentration of the solitary receptors changed only moderately across the convex hull (Fig. 4b). Thus, the solitary receptors appear to diffuse more readily into and out of the mesophasic assembly.

**Fig. 4.**
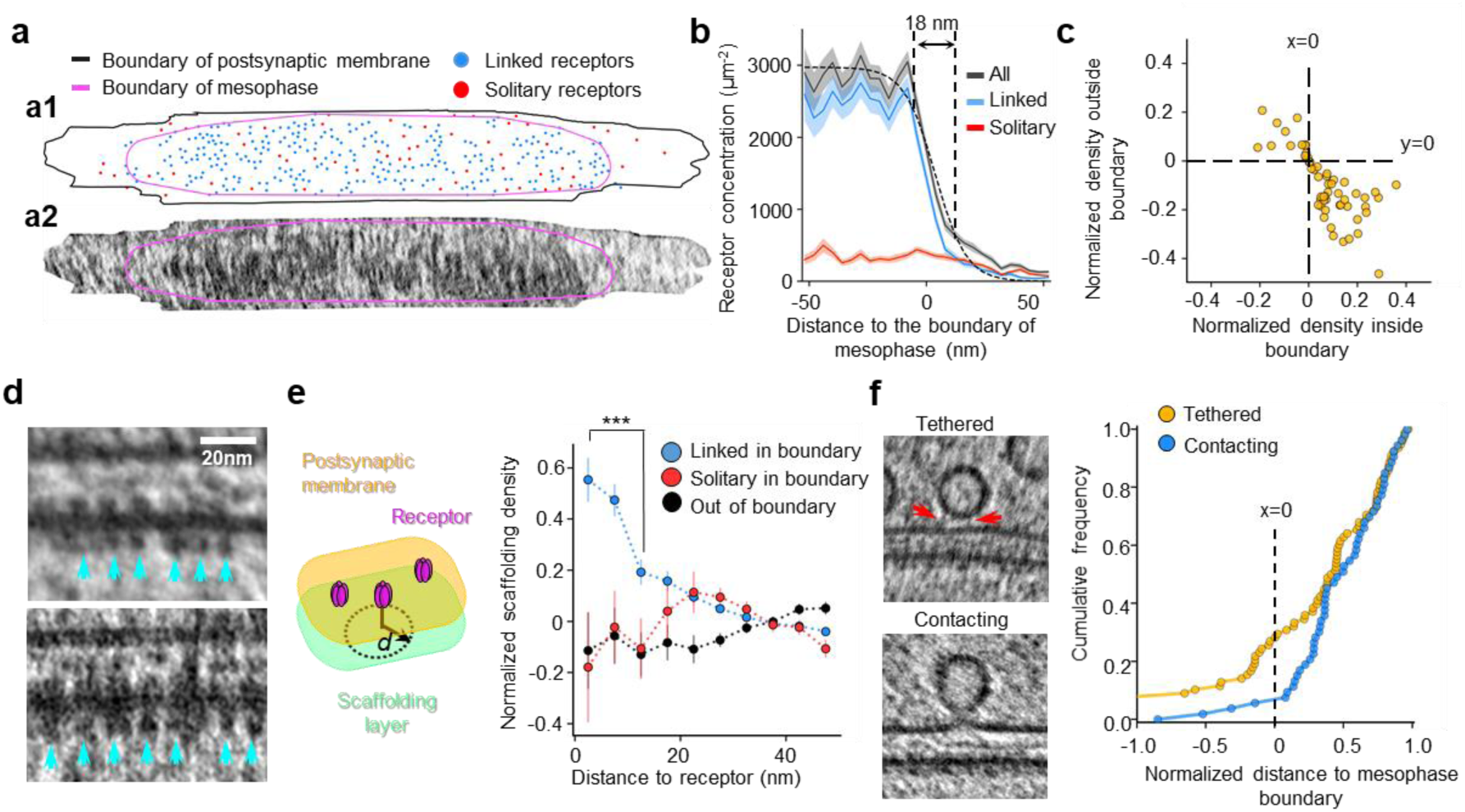
Mesophasic assembly of inhibitory PSD. **a,** Examples of receptor distribution on the postsynaptic membrane (**a1**) and the corresponding density projection of the scaffolding layer (**a2**). **b,** Receptor concentration as a function of distance to the mesophase boundary (n=58). The dashed curve is a sigmoid function fitted with the black curve. Light shadow: SEM. Vertical dashed lines: 80%-20% width of the sigmoid function. **c,** Correlation of scaffolding layer density inside and outside mesophase. **d,** Example of interactions between receptors and scaffolding proteins. Cyan arrows indicate density of scaffolding proteins interacting with GABA_A_Rs. **e,** Left: Diagram showing relative positions of scaffolding layer and receptors, with *d* representing distance to the projection of receptor in scaffolding layer. Right: Normalized scaffolding density as a function of *d* (n=9,531). ***P<0.001, two-tailed t-test. **f,** Left: Example tomographic slices of tethered and contacting vesicles. Red arrows indicate rod-like tethers. Right: Cumulative frequency of normalized distance from vesicles to mesophase boundary. A vesicle with normalized distance of 1 means it is at the center of the mesophasic condensate, while a vesicle with normalized distance of 0 means it is on the mesophase boundary. N=81 for tethered vesicles; N=54 for contacting vesicles. The distributions of the two vesicle populations are significantly different (P=0.013, two-tailed t-test).

It is known that GABA_A_R interacts with scaffolding molecules gephyrin and associated proteins that may interact with one another to form thin sheet-like densities in parallel to the postsynaptic membrane^7^. To examine whether such interactions might underlie the observed organization of GABA_A_Rs, we obtained a 2D density projection of the scaffolding layer (Fig. 4a2). Distinct condensate-like densities were observed in the scaffolding layer (Fig. 4a2; Extended Data Fig. 7b), well correlated with the mesophasic assembly of GABA_A_Rs in majority of synapses (Fig. 4c). Furthermore, many particles in the scaffolding layer positioned directly underneath individual receptor densities, also with ∼11 nm inter-particle distances (Fig. 4d). Quantitative analysis further confirmed that the density in the scaffolding layer was higher directly underneath a linked GABA_A_R within the phase boundary (Fig. 4e). In contrast, the higher peri-receptor scaffolding density was not observed for receptors outside the phase boundary, nor was it found for solitary receptors within the phase boundary, indicating that such receptors might not have direct interactions with the scaffolding molecules (Fig. 4e). Thus, the semi-ordered organization of linked receptors is likely due to their interaction with the underlying scaffolding molecules, which form a semi-ordered sheet-like condensate, probably through multivalent interactions.

### Mesophasic organization of PSD correlates with neurotransmitter release

It is tempting to hypothesize that the mesophasic organization of GABA_A_Rs may functionally correlate with presynaptic neurotransmitter release. To test this, we analyzed the locations of synaptic vesicles near the presynaptic active zone. In our tomograms, two types of vesicles were identified: one tethered to the presynaptic membrane through rod-like densities, thus termed hereafter as tethered vesicles; the other had direct contact with the presynaptic membrane, thus termed as contacting vesicle (Fig. 4f). Both types of vesicle-plasma membrane interaction have also been observed in cryoET studies of purified synaptosomes^36^. Intriguingly, most (93%) of the contacting vesicles located within the presynaptic area apposing to the postsynaptic region inside the phase boundary (Fig. 4f; Extended Data Fig. 7d). Outside the boundary, the number of contacting vesicles are significantly fewer as compared to that expected from random distribution. In contrast, the number of tethered vesicles located inside or outside this area are not significantly different from that of expected from random distribution (Fig. 4f; Extended Data Fig. 7d). It has been suggested that tethering allows initial targeting of vesicles to the membrane, and the contacting vesicles are more ready to release upon stimulation^36,37^. If this is the case, our observations suggest that vesicular GABA is primarily released towards the semi-ordered GABA_A_R networks within the mesophasic boundary, thus optimizing the efficiency of neurotransmission.

## Discussion

By quantitative analysis of individual GABA_A_Rs in intact inhibitory synapses, our results reveal a unique mesophasic state of receptor assembly that is likely to result from the binding of these receptors to mutually-interacting scaffolding proteins such as gephyrin and their associated proteins^38,39^. Unlike the previous picture of a hexagonal lattice-shaped PSD architecture^38,40,41^, we observed semi-ordered receptor networks with a fixed value, 11-nm, for preferred inter-receptors distance, and with virtually uniformly distributed values for relative angles. This unique property suggests that the interaction among gephyrin molecules and associated proteins are also semi-ordered, forming flexible networks. Importantly, this network is confined to a two-dimensional sheet parallel to the postsynaptic membrane^7^, probably by the interaction of gephyrin with membrane-bound GABA_A_Rs. Such a mesophasic assembly exhibits both variability and regularity, demonstrating how ensembles of synaptic molecules acquire great complexity via self-organization. This organization principle may also suggest a molecular strategy for a synapse to achieve its “Goldilocks” state with a delicate balance between stability and flexibility on the micro-nano scale.

## End notes

### Supplementary information

Supplementary Information is linked to the online version of the paper.

## Acknowledgments

We thank Dr. Peng Ge for technical advice on cryoEM imaging and Dr. Aihui Tang for valuable suggestions on the manuscript. This work was supported in part by grants from the Strategic Priority Research Program of the Chinese Academy of Sciences (XDB32030200), the National Natural Science Foundation of China (31630030, 31621002, 31761163006, and 31600606), the National Key R&D Program of China (2017YFA0505300 and 2016YFA0501100), the China Postdoctoral Science Foundation (2018M640590), and the Anhui Provincial Natural Science Foundation (1908085QC95). Research in the Zhou group is supported in part by the U.S. National Institutes of Health (GM071940). We acknowledge use of instruments at the Center for Integrative Imaging of Hefei National Laboratory for Physical Sciences at the Microscale, and those at the Electron Imaging Center for Nanomachines of UCLA supported by U.S. NIH (S10RR23057 and S10OD018111) and U.S. NSF (DMR-1548924 and DBI-133813). We thank the Bioinformatics Center of the University of Science and Technology of China, School of Life Sciences, for providing supercomputing resources for this project.

## Author contributions

Y.-T.L., C.-L.T., P.-M.L., Z.H.Z., and G.-Q.B. designed research; C.-L.T., R.S., and L.Q. performed experiments; Y.-T.L., C.-L.T., X. Z., and G.-Q.B. analyzed data; Y.-T.L., C.-L.T., P.-M.L., Z.H.Z., and G.-Q.B. wrote the paper. All the authors edited and approved the manuscript.

The authors declare no competing financial interests.

## Data and materials availability

All data reported in the main text or the supplementary materials are available upon request.

## Code availability

All code used in the paper is available upon request.

## Methods

All animal experiments were approved by the Animal Experiments Committee at the University of Science and Technology of China.

### Primary culture of hippocampal neurons

Low-density cultures of dissociated embryonic rat hippocampal neurons were prepared according to the protocols described previously^7^. In brief, electron microscopy (EM) gold finder grids (Quantifoil R2/2 Au NH2 grids) were plasma cleaned with H_2_ and O_2_ for 10 s using a plasma cleaning system (Gatan), sterilized with UV light for 30 min, and then treated with poly-L-lysine before use. Hippocampi were dissected from embryonic day-18 rats and were treated with trypsin for 15 min at 37 °C. The dissociated cells were plated on the treated EM grids at a density of 40,000-60,000 cells/ml in 35-mm Petri-dish, and maintained in incubators at 37 °C in 5% CO_2_ atmosphere. NeuroBasal (NB) medium (Invitrogen) supplemented with 5% heat-inactivated bovine calf serum (PAA), 5% heat-inactivated fetal bovine serum (HyClone), 1× Glutamax (Invitrogen) and 1× B27 (Invitrogen) was used as culture medium. Each Petri-dish was added with 1.5 ml medium. Twenty-four hours after plating, half of the medium was replaced with serum-free culture medium. Then, one-third of the culture medium was replaced with fresh serum-free culture medium every three days. The cultures were treated with cytosine arabinoside (Sigma) to prevent the overgrow of glia cells. Some of the cultures experienced inactivation by 2-day treatment with 1 μM tetrodotoxin (TTX) or 1-hour treatment with 2 μM TTX followed by 3-hour treatment with 2 μM TTX plus 50 μM APV^42,43^. We did not observe significant difference among different groups in basic properties of receptor expression and organization and thus pooled the data together for all analyses. For cryo correlative light and electron microscopy (cryoCLEM) experiments, cultures were transfected with mCherry-gephyrin (a gift from Dr. Ann Marie Craig) using Lentivirus at 10 DIV, as described previously^7^. All cultures were used for cryo-electron tomography (cryoET) imaging at 16 days *in vitro* (DIV).

### Frozen-hydrated sample preparation

At DIV 16, the culture medium was replaced with extracellular solution (ECS, containing 150 mM NaCl, 3 mM KCl, 3 mM CaCl_2_, 2 mM MgCl_2_, 10 mM HEPES and 5 mM glucose, pH 7.3). The EM grids were taken out from the CO_2_ incubator and loaded into a Vitrobot IV (FEI) which was maintained in 100% humidity. Protein A-coated colloidal gold beads (15-nm size, CMC) were added to the grid (4 μl each, stock solution washed in ECS and diluted 10 times after centrifugation) as fiducial markers. The grids were then blotted and plunged into liquid ethane and were stored in liquid nitrogen until use.

### CryoCLEM imaging

For cryoCLEM imaging, we used the same procedures in our previous paper^7,44^. In brief, the inside channel of the custom built cryo-chamber was precooled to −190 °C by liquid nitrogen, and maintained below −180 °C. Then, an EM grid with frozen-hydrated sample was loaded onto an EM cryo-holder (GATAN), which was subsequently inserted into the cryo-chamber. Dry nitrogen gas flowed around the 40X objective lens (Olympus LUCPLFLN 40×, NA 0.6) throughout the experiment to prevent the formation of frost. Fluorescence images were taken with an ANDOR NEO sCMOS camera (Andor) attached to the fluorescence microscope. For each field of view, both bright-field and mCherry channel (Ex: 562/40, DM: 593, Em: 641/75; Semrock, mCherry-B-000) images were acquired.

The EM cryo-holder with the grid was then directly transferred into a Tecnai F20 microscope (Thermo Fisher). Indexes of the finder grids were used to roughly identify the areas of the sample imaged in cryo-light microscope. Then *Midas* program in IMOD package^45^ was used to roughly align the low magnification (330×) EM images with the bright-field LM images. After rough alignment, a set of holes (about 10 for each images) on the carbon film of the grid were manually picked using 3dmod in IMOD package from both the low magnification EM images and their corresponding fluorescence images. Transformation functions between the EM and LM images was calculated by correlating the selected positions in both images.

After aligning the low magnification EM images with LM images, pixel-wise positions of ∼15 holes on carbon film (in one square) in each low magnification EM image were recorded. Afterwards, those holes were identified at 5,000× magnification and their mechanical coordinates (*i.e.* positions on the EM sample stage) were also recorded. The transformation function from the pixel-wise positions to EM mechanical coordinates was determined. Then the puncta of gephyrin-mCherry were selected manually using 3dmod in IMOD. Positions of these fluorescent puncta were then converted into corresponding EM mechanical coordinates with the transformation functions to guide tilt series acquisition. Finally, reconstructed tomographic slices were fine-aligned and merged with the fluorescence images to identify each synapse (Fig. 1a) using Midas and ImageJ.

### Cryo-electron tomography (CryoET) imaging

For cryoCLEM experiments, the tilt series were collected using a Tecnai F20 microscope (Thermo Fisher) equipped with Eagle CCD camera (Thermo Fisher). The Tecnai F20 was operated at an acceleration voltage of 200 kV. Tilt series were collected first from 0° to −60° and then from +2° to +60° at 2° intervals using FEI Xplore 3D software, with the defocus value set at −12 to −18 µm, and the total electron dosage of about 100 e^-^/Å^2^. The final pixel size was 0.755 nm.

For the analysis of GABA_A_Rs, cryoET data were collected using a Titan Krios (Thermo Fisher) equipped with a Volta phase plate (VPP), a post-column energy filter (Gatan image filter), and a K2 Summit direct electron detector (Gatan). The energy filter slit was set at 20 eV. The Titan Krios was operated at an acceleration voltage of 300 KV. When VPP was used, defocus value was maintained at −1 μm; otherwise, it was maintained at −4 μm. The VPP was conditioned by pre-irradiation for 60 s to achieve an initial phase shift of about 0.3π. Images were collected by the K2 camera in counting mode or super-resolution mode. When counting mode was used, the pixel size was 0.435 nm. For super-resolution mode image, the final pixel size was 0.265 nm. Tilt series were acquired using SerialEM^46^ with two tilt schemes: from +48° to −60° and from +50° to +66° at an interval of 2°; from +48° to −60° and from +51° to +66° at an interval of 3°. The total accumulated dose is ∼150 e^-^/Å^2^. For sub-tomogram analysis, 6 grids were used for data collection. Totally, 32 and 40 inhibitory synapses were imaged with and without VPP, respectively.

### 3D reconstruction of the tomograms

Each recorded movie stack was drift-corrected and averaged to produce a corresponding micrograph using MotionCorr^47^. To combine the data with different pixel size during image processing, we rescaled the images recorded with super-resolution mode with antialiasing filter to match the pixel size of image recorded with counting mode (0.435 nm/pixel), by *newstack* command in IMOD. For images recorded without VPP, the defocus value of each image was determined by CTFFIND4^48^. For tilt series acquired with VPP, the defocus values cannot be precisely calculated. However, the defocus of each image is relatively low (∼1 um), which does not limit the resolution obtained by sub-tomogram averaging. Thus, we did not perform defocus determination and CTF correction for these tilt series.

Tilt series were aligned with 15 nm gold beads as fiducial markers using IMOD. 3D reconstruction was performed with weighted back projection algorithm (WBP) using NovaCTF^49^. Because those tomograms had low contrast and were hard to interpret by visual inspection, we also used SIRT-like filter in NovaCTF to generate tomograms equivalent to those reconstructed by SIRT algorithm with 5 iterations. Segmentation and cryoET density analyses were performed using the SIRT-like filter reconstructed tomogram, whereas sub-tomogram averaging was performed using tomogram reconstructed with WBP.

Because the samples are thick, to eliminate the depth-of-the-focus problem, we performed 3D-CTF correction^49^ and obtained CTF phase flipped tomograms for tilt series acquired without VPP. The defocus step for depth-of-the-focus correction was 50 nm.

### 3D rendering

By manually placing markers corresponding structures using *volume tracer* in UCSF Chimera^50^, synaptic membranes and organelles such as microtubules, actin filaments, mitochondria and multivesicular bodies were traced and segmented. Then, the manually segmented structures were smoothed by Gaussian filter. The ribosomes and synaptic vesicles were identified by template matching using PyTom^51^, as described previously^7^. The vesicles were rendered based on their diameter.

### Generating uniform oversampled points on postsynaptic membranes

Previous study showed the sub-tomogram average can be preformed with uniform selected sub-tomograms on a given surface taking advantage of the geometry of that surface^52,53^. We thus sought to reconstruct the structure of GABA_A_R by uniform oversampled sub-tomogram on the postsynaptic membrane segmented manually. Postsynaptic membrane was defined as the synaptic membrane area corresponding to the uniform synaptic cleft.

To segment postsynaptic membrane, we first segmented the synaptic cleft volumes in two-times binned tomograms using segmentation tool in Amira (Thermo Fisher). As the pixel size of all tomogram was or was scaled to 0.435 nm/pixel, the pixel size of two-times binned tomograms was 0.87 nm/pixel. Then we used Sobel filter to generate boundary surface of the segmented synaptic cleft. This boundary represented two opposed membranes: presynaptic and postsynaptic membrane. Then the postsynaptic membrane was manually extracted.

To generate uniformly oversampled points, we first generated a uniformly distributed 3D lattice of hexagonal close-packaging points in two-times binned tomograms (Extended Data Fig. 1a). The distance between the two nearest sampling points in the lattice is 5 pixels (4.35 nm). All the sampling points were within 8.7 nm distance to the segmented membrane. The two-times binned sub-tomograms, whose centers are the sampling points, were then extracted using *boxstartend* program in IMOD. The extracted box size of each sub-tomogram is 32×32×32 pixels (27.84×27.84×27.84 nm). Because the sampling distance is 5 pixels, the nearest distance from the center of any possible receptor to the one sampling point is less than 2.5 pixels. Given 7 nm (∼8 pixels) diameter of GABA_A_R, each receptor should be fully covered in multiple extracted sub-tomograms so that no receptor was omitted during sampling.

The orientation of each sub-tomograms has three Euler angles denoted as parameters within the Relion star file^25^: rot (_rlnAngleRot), tilt (_rlnAngleTilt) and psi (_rlnAnglePsi). During the sub-tomogram extraction, the initial tilt and psi angles of each sub-tomogram were calculated as the orientation perpendicular to the patch of membrane in that sub-tomogram. The rot angle (rotational angle around the vector that is perpendicular to the membrane) for each sub-tomogram was set randomly.

With the uniform oversampling, we obtained 171,374 and 135,717 two-times binned sub-tomograms near postsynaptic membrane from tomograms imaged with and without VPP, respectively.

### Initial 3D classification using two-times binned sub-tomograms

The classification and refinement of the sub-tomograms were preformed using Relion (Extended Data Fig. 1b)^25,54^. The tomograms imaged with and without VPP appeared to be with different contrast. It is possible that 3D classification classifies the same protein feature into different classes based on whether the sub-tomogram was acquired with VPP or not. To minimize this error of classification, we performed the classification separately for sub-tomograms imaged with VPP and without VPP. This separation also enables cross-validation between results obtained from data acquired with VPP and without VPP (Extended Data Fig. 1b).

To identify GABA_A_Rs containing tomograms from those sub-tomograms, we performed 3D classification imposing 5-fold symmetry using Relion3. The resolution for the classifications was limited to 30 Å. To ensure that the orientation was searched around the vector perpendicular to membrane, we set the prior of tilt and psi angles as the calculated angles corresponding to the orientation of membrane and set the sigma of local angle search for tilt and psi angles as 3 degrees. We did not set any limitation in searching for the rot angle during classification. To limit the 3D positional search during 3D classification, the prior of the offset searching range was set as 0, meaning the offset was only searched around the center of the sub-tomograms. The offset search range was set to ±3 pixels. The initial reference was generated by *relion_reconstruct* using the predetermined Euler angles. As expected, the initial reference appeared as a flat membrane structure due to the averaging of uniform oversampled sub-tomograms on the membrane (Extended Data Fig. 1c). Because tilt series imaged without VPP were corrected using 3D-CTF and the tilt series imaged with VPP were recorded at low defocus value (−1 um), we did not perform CTF correction during image processing using Relion. To compensate missing wedge, missing wedge volumes (_rlnCTFimage in relion star file), which were 3D masks in Fourier space, were generated by custom made scripts. The classifications were performed with 100 iterations (Extended Data Fig. 1c).

To determine the optimal number of classes for 3D classification, we tested the number of classes from 8 to 15 in classification. We obtained one ‘good’ class, which appeared similar to previously published GABA_A_R structures, for all number of classes from 8 to 13 during the classification. The number of sub-tomograms in the ‘good’ class reduced as the number of classes increased from 8 to 11 but became stable after 11 (Extended Data Fig. 2a). The structures of the classification result became worse when the number of class was larger than 13. Thus, we used 12 as the optimal number of classes for classification and obtained the ‘good’ class among the 12 classes for both of the classifications using data collected with and without VPP (Extended Data Fig. 1b).

To eliminate that two or more sub-tomograms corresponding to the same receptor, we removed duplicated sub-tomograms as follows. We mapped the refined positions of the sub-tomograms after 3D classification to the original tomograms. If distances between centers of two classified sub-tomograms in original tomogram were smaller than 7 nm (the diameter of GABA_A_R), the sub-tomogram with lower score (_rlnLogLikelihoodContribution in Relion star file) were removed. After removing duplicate, we obtained 7,089 and 5,004 sub-tomograms obtained from data acquired with and without VPP, respectively.

### First round of 3D refinement using unbinned sub-tomograms

Then we calculated the coordinates of sub-tomograms in the corresponding unbinned original tomograms (with pixel size of 4.35 Å/pixel) and extracted new sub-tomograms with box size of 64×64×64 pixels. We combined sub-tomograms from VPP and no VPP data for 3D auto-refine (Extended Data Fig. 1b). Then, we generated 60 Å resolution initial references by *relion_resonstruct* with the predetermined orientations. Differing from the previous round of classification, we didn’t limit the search angle and didn’t set prior for angle and offset searching during the 3D refinement. Five-fold symmetry was imposed during 3D refinement. This round of 3D auto-refine refined the orientation and the positions of sub-tomograms and generated a preliminary reconstruction at 21 Å resolution, which was reported during *relion_refine* processing. The duplicated sub-tomograms were further removed. After this step, we obtained 6,919 and 4,904 sub-tomograms for VPP and no VPP data, respectively.

### Removing outliers of tilt and psi angles

Because synaptic membrane is relatively flat and GABA_A_Rs are perpendicular to the membrane, the tilt and psi angles for sub-tomograms should be similar in each synapse. Thus, we used this knowledge to further reduce the error of receptor identification, as follows. We plotted the distributions of tilt and psi angles for the sub-tomograms in each synapse (Extended Data Figs. 2b-c). Indeed, the distribution of the refined tilt and psi angles of sub-tomograms in a given synapse were in a cluster with approximately Gaussian distribution, whose center corresponds to the angles perpendicular to the postsynaptic membrane (Extended Data Fig. 2c). Few sub-tomograms have orientations perpendicular to the membrane but pointing to the cytoplasmic side, possibly they were aligned to the proteins of postsynaptic densities (PSD) on the cytoplasmic side. We discarded those sub-tomograms for further refinement. The percentage of those misaligned sub-tomograms with opposite orientation are 2% and 5% for VPP and no VPP data, respectively (Extended Data Fig. 1d). Furthermore, we also excluded sub-tomograms whose tilt and psi angles are three times of standard deviation (σ) away from the center of the Gaussian distribution (10% and 13% of total sub-tomograms from VPP and no VPP data, respectively) (Extended Data Figs. 2c-e).

### Removing outliers of low score

Then, we removed sub-tomograms with lower scores (_rlnLogLikelihoodContribution in Relion star file). We normalized the scores of sub-tomograms in each synapse, ensuring the normalized scores of the sub-tomograms for each synapse has an average of 0 and a standard deviation of 1. The distribution of normalized scores is a slightly lopsided Gaussian distribution (Extended Data Fig. 2f). We fitted the distribution with a Gaussian distribution and then removed the sub-tomograms with scores less than mean minus 2σ. The ratio of sub-tomograms with lower score were ∼3% for both VPP and no VPP data (Extended Data Figs. 2e-g).

### Second round of 3D refinement using unbinned sub-tomograms

After removing outliers, those sub-tomograms were used for a second round of 3D auto-refine (Extended Data Fig. 1b). Local searches with sigma angle of 3° for orientation determination were performed during 3D auto-refinement. Five-fold symmetry was imposed during 3D refinement. The final resolution of the reconstruction was estimated with two independently refined maps from halves of the dataset with gold-standard Fourier shell correlation (FSC) at the 0.143 criterion^55^ using *relion_postprocess*, and was determined to be 19 Å (Fig. 1g).

### Analysis the accuracy of rot angle

To estimate the accuracy of rot angle, we calculated two sets of cross-correlation (CC) score for the original sub-tomograms and sub-tomograms that rotated 36⁰ (Extended Data Fig. 2h). CC score represents the similarity between a sub-tomogram and the sub-tomogram average of GABA_A_R. To do so, we rotated the sub-tomogram average by 36⁰, and then processed the sub-tomograms with *relion_refine* using original and rotated sub-tomogram averages as references, separately. We skipped both maximization step and alignment step in order to prevent updating references and orientation search, respectively. We used *always_cc* argument to calculate the CC score instead of log likelihood that was default in Relion. The processes were finalized with 1 interaction. By this processing, we obtained two new star files with the CC scores. We plotted the distribution of CC scores in the two star files. Indeed, the score distributions for the two sets of sub-tomograms are well separated (Extended Data Figs. 2h-i).

### Estimate error rate of receptor identification

To estimate the error of our 3D classification with uniformly over-sampled sub-tomograms, we visually inspected all the identified receptors in 4 selected tomograms acquired with VPP. Few receptors identified by our methods cannot be recognized, thus those receptors could be falsely identified. Thus, the error rate was defined as the percentage of identified receptor that cannot be recognized visually for each synapse.

The error rates for the 4 synapses are 14.4% (16 out of 111), 6.0% (5 out of 83), 22.9% (32 out of 140), and 18.3% (62 out of 339), respectively.

### False positive rate of receptor identification

In order to evaluate the false positive rate of GABA_A_R identification, we repeated the sub-tomogram analysis using data mixing the same sub-tomograms and intentionally induced negative controlled sub-tomograms on presynaptic membrane (Extended Data Fig. 3). These negative controlled sub-tomograms were extracted using the same uniform oversampling methods on the segmented presynaptic membranes. Presynaptic membranes of 2 inhibitory synapses imaged with VPP and 2 inhibitory synapses imaged without VPP were used for this analysis.

We did the classifications and refinements (Extended Data Figs. 3a-b) exactly the same as the previous described steps. The classifications and refinements with data mixing with negative controlled sub-tomograms also generated structures of GABA_A_Rs. As expected, the number of GABA_A_Rs identified using sub-tomograms with negative controlled sub-tomograms for each synapse is similar to the receptor identified without negative controlled sub-tomograms (Extended Data Fig. 3c).

For synapses analyzed for both pre- and post- synaptic membrane, we calculated false positive rate as falsely identified receptors on presynaptic membranes dividing number of receptors on post synaptic membranes. The false positive rates for the two synapses imaged with VPP are 15% and 10%. The false positive rates for the two synapses imaged without VPP are 13% and 10% (Extended Data Fig. 3d).

### 3D classification of the oversampled sub-tomograms without symmetry

We then tested whether the classification without symmetry could yield structures similar to the GABA_A_R structure published before. We used the same sub-tomograms acquired with VPP and performed the classification without symmetry. The other parameters were the same as the first round classification described before. Indeed, this classification generated structures with sizes similar to the GABA_A_R. However, the structures were worse than the reconstruction with 5-fold symmetry and were not centered properly (Extended Data Fig. 4a). Intriguingly, two receptor-like structures could present in the same sub-tomogram average (Extended Data Fig. 4b). This further confirmed that the receptors tend to form receptor pairs with 11 nm inter-receptor distance.

### Analysis and reconstruction of receptor pair

For each receptor pair (with 11±4 nm inter-particles distance), we calculated the coordinate of the center of the two GABA_A_Rs, and used this coordinate to extract sub-tomograms (64×64×64 pixels) in two-times binned original tomograms. The tilt and psi angles of a receptor pair was set as the mean of those angles for the two receptors. The rot angle was calculated to ensure that the vector from one receptor to the other receptor aligns to the x axis of the receptor pair reconstruction (Fig. 2f). We then reconstructed receptor pair using *relion_reconstruct* with the calculated orientations. Total 15,184 sub-tomograms of receptor pairs were used in the reconstruction.

### Measuring the angle (ω) between the rotation of the receptor and pair axis

We then calculated the angle (ω) between the rotation angle (rot) of one given receptor in a receptor pair to receptor pair axis (Fig. 2g). The receptor pair axis was defined as a vector from the other receptor to the given receptor. Then we separated the sub-tomograms into four groups by the ω angle— 0-18°, 18-36°, 36-54°, 54-72° groups, containing 3,883; 2,957; 4,199 and 4,195 sub-tomograms, respectively. We further reconstructed the sub-tomograms in each group using *relion_reconstruct*. In all four reconstructions, the given receptor appears to have pseudo 5-fold symmetry.

### Reconstructing receptor triplet and analyzing the angle θ between the two arms

One GABA_A_R can also pair with two neighbor receptors forming receptor triplet. Each triplet has two arms, which connect the central receptor to the two neighbor receptors. We then calculated the angle θ between the two arms of the triplet (Fig. 2i). We reconstructed the triplets with 50-70°, 80-100°, 110-130°, 160-180° of θ value, containing 2,428; 1,772; 1,937 and 1,714 sub-tomograms, respectively (Fig. 2j). The center of each receptor triplet sub-tomogram was set as the mass center of the three receptors. Tilt and psi angles of each sub-tomogram were set as the mean angles of the three receptors. The rot angle of a receptor triplet sub-tomogram was calculated to ensure that the vector from one neighbor receptor to the other is parallel to x axis. All reconstructions were computed using *relion_reconstruct* by the sub-tomograms (64×64×64 pixels) extracted from two-times binned tomograms.

### Local receptor concentration and nearest neighbor distance analysis

Among the 72 synapses we obtained, 2 of them imaged without VPP were not fully covered in the tomograms. These two synapses were excluded in the analyses of GABA_A_Rs distribution in the following sections.

We calculated the concentration of receptors around a given point on the membrane. In our case, the given point is either a receptor or random selected point on postsynaptic membrane. We partitioned the membrane around the given point into concentric rings of 2 nm width. The radius range of the rings is from 0 to 32 nm. Then, the receptor concentration was calculated as number of receptors in a ring dividing the surface area of that ring.

We also calculated the first and second nearest neighbor distance for each receptor, using standard distance formula in 3D.

### Analysis of the receptor networks

If two receptors have distance smaller than 15 nm, they were defined as “linked” receptors. We then defined a receptor network as follows. If two receptors are linked by series of (equal or more than 0) receptors, we grouped them in the same network. Otherwise, they are in different networks. The network size was defined as number of receptors in a network. Randomized receptor distributions were generated from the same number of receptors over the same postsynaptic area.

### Calculation of the Voronoi entropy

To calculate the Voronoi entropy of each synapse, we first calculated the first two principle vectors for all 3D segmented points on postsynaptic membrane, using singular value decomposition in Matlab. Using the two principle vectors, we projected the 3D receptor locations on a 2D plane. Then we generated Voronoi tessellation of the 2D locations of receptors in each synapse (Extended Data Fig. 6c) using scipy.spatial.Voronoi function in SciPy (https://scipy.org). Voronoi entropy was calculated using follow formula^32^:

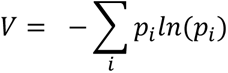

Where *i* is the number of vertices of a polygon. *p_i_* is the frequency of the polygon with *i* vertices. *ln* is the natural logarithm. *V* is the Voronoi entropy.

### Determining the boundary of mesophasic assembly of GABA_A_Rs

To determine the boundary of the receptor assembly, the receptor positions were first projected onto the 2D plane as described before. Then, a convex hull of all linked receptors for each synapse were constructed using python package *shapely* (https://github.com/Toblerity/Shapely). To eliminated the coincidently formed linked receptors outside the condensed receptors region, we smoothed the convex hull by 40 nm dilation followed by 40 nm erosion using python package shapely (Extended Data Fig. 7a). Convex hull of 12 (out of 70) synapses have diameter smaller than 80 nm. Those synapses are not eligible for dilation, so they were excluded in the phase boundary analysis. The distance of a receptor to mesophase boundary was also calculated using shapely.

### Calculation of synaptic membrane area

To calculate the area of postsynaptic membrane, we first generated surface of the postsynaptic membrane in 3D using *imodmesh* in IMOD. The area of postsynaptic membrane was extracted from output of *imodinfo* command in IMOD. While in Fig. 4b and Extended Data Fig. 7c, the postsynaptic membranes were projected to a two dimensional plane. Thus in those figures, membrane areas were calculated two dimensionally using *shapely*.

### Analysis electron microscopy density of scaffolding layer

To analysis the density of scaffolding layer, we first extracted the voxels in the scaffolding layer region in the tomogram as densities 10-15 nm toward the cytoplasmic side from the postsynaptic membrane. The scaffolding layer region were then project to a 2D plane, using two principle vectors of postsynaptic membrane described previously, resulting in 2D density profiles of the scaffolding layer parallel to the postsynaptic membrane. The 2D density profiles were then normalized so that the mean pixel density of the profiles is 0 and the standard deviation is 1. The mesophase boundary of GABA_A_R were mapped on the 2D profile of the scaffolding layer. Densities inside and outside mesophase boundary on the 2D profiles were calculated as the mean pixel density inside and outside of the boundary, respectively.

We then calculated the density of scaffolding layer around 2D projected locations of receptors (Fig. 4e). We partitioned the 2D profile of scaffolding layer around a receptor into concentric rings of 5 nm width. The radius range of the rings is from 0 to 50 nm. Then, the density of scaffolding layer was calculated as the mean intensity value both in the concentric ring and inside postsynaptic membrane area (Fig. 4a). Hence, we produced the relation between the distance to the given receptor and the pixel density values of the 2D profile of scaffolding layer.

### Analysis of tethered and contacting synaptic vesicles

To calculate distance from synaptic vesicle to mesophase boundary, we first manually selected positions on the presynaptic membrane nearest to a contacting or a tethered vesicle. Then, the positions were projected to 2D plane using the methods described before. The distance (d1) from the synaptic vesicle projection point to mesophase boundary were calculated using *shapely*. We also project all segmented points inside mesophase on postsynaptic membrane to the 2D plane and calculated the largest distance (d2) from those points to the mesophase boundary. The normalized distance from synaptic vesicle to mesophase boundary was calculated as d1/d2. Randomized vesicles were generated by randomly selecting locations over the same synaptic area. We repeated the randomization 10 times for each synapses. The mean number of randomized vesicles inside or outside of mesophase boundary were used for statistical analysis.

## Extended data figures, table, and legends

**Extended Data Fig. 1.**
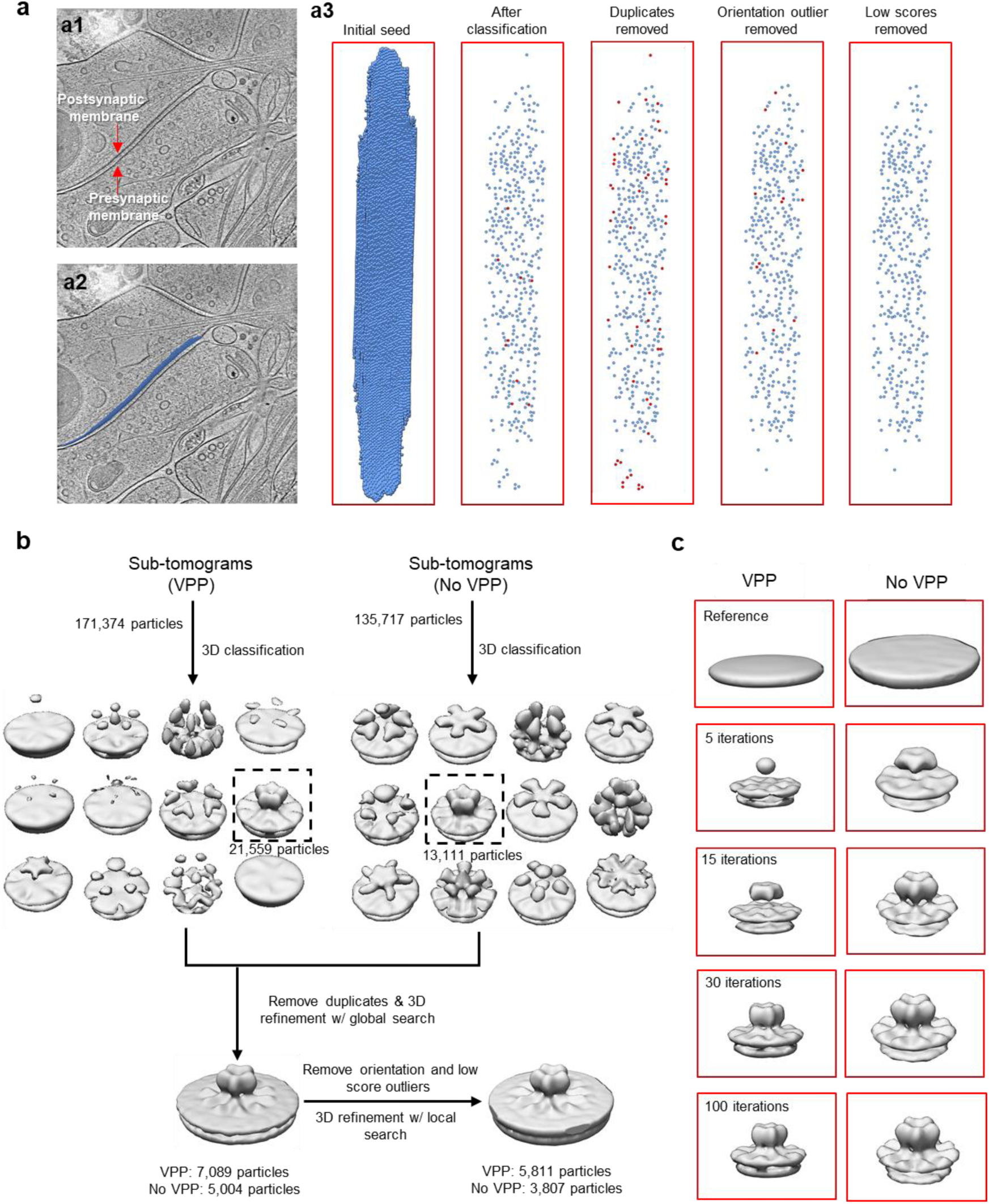
Flowchart illustrates identification and sub-tomogram averaging of GABA_A_R. **a,** Steps for identifying GABA_A_Rs from sampling points. **a1,** Electron tomographic slice of an inhibitory synapse. **a2,** Electron tomographic slice superposed with sampling points on postsynaptic membrane. **a3,** Sampling points after each step. Red points are sampling points that will be discarded in the next step. **b,** Classification and refinement of GABA_A_R on postsynaptic membrane. **c,** Structure of GABA_A_R emerged during iterative classification.

**Extended Data Fig. 2.**
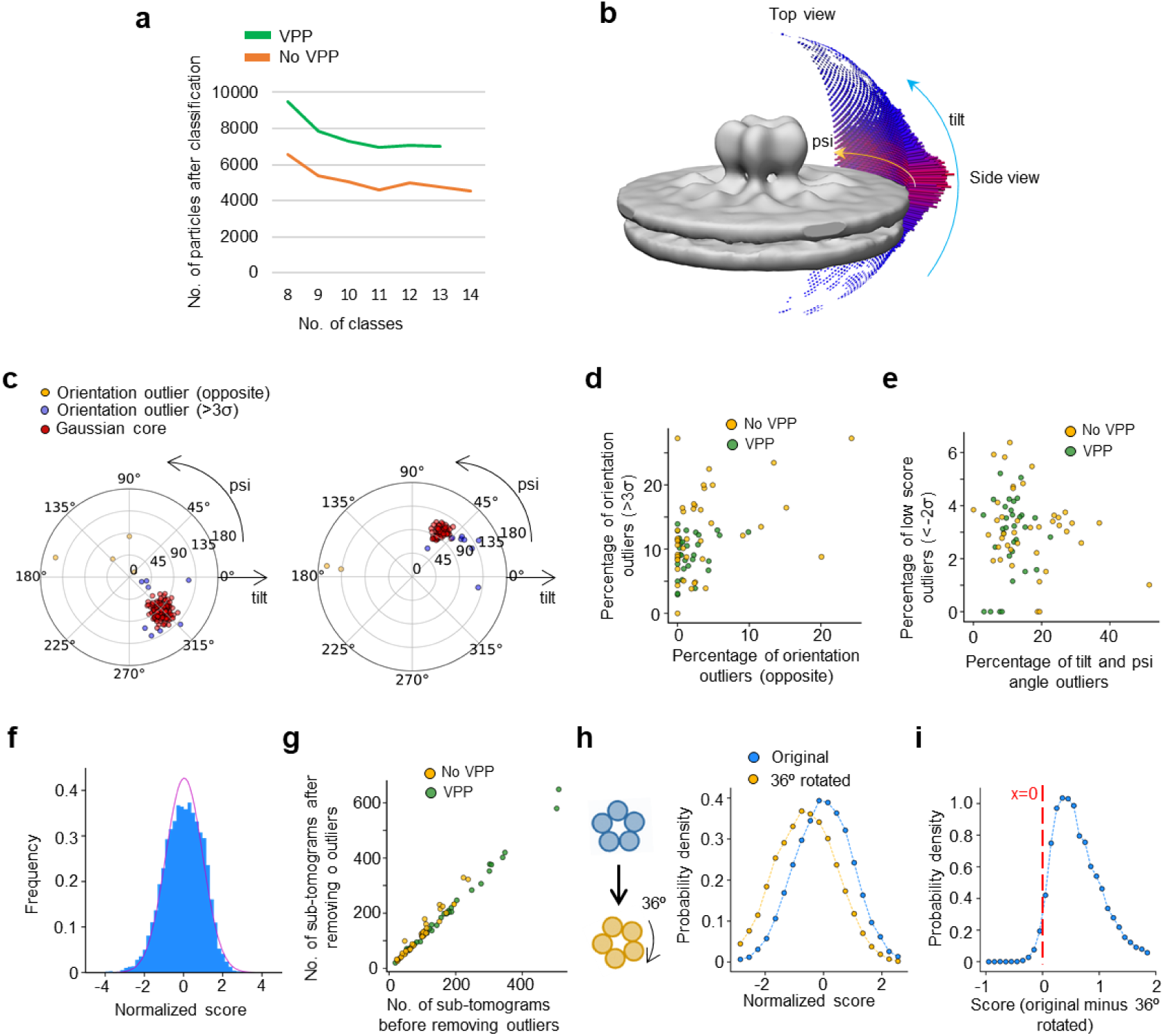
Performance estimation of template-free classification and refinement. **a,** Number of classified GABA_A_R sub-tomograms plotted against number of classes. **b,** Euler (psi and tilt) angles of all sub-tomograms used for the final sub-tomogram averaging. **c,** Distribution of Euler angles for sub-tomograms in two example synapses after first round of refinement. **d,** Percentage of outliers with opposite angles verses percentage of outliers with angles 3σ away from Gaussian core in each synapse. **e,** Percentage of all orientation outliers verses percentage of low score outliers in each synapse. **f,** Frequency distribution of sub-tomogram scores fitted with Gaussian curve (red curve). **g,** Number of sub-tomograms before and after removing both orientation and low score outliers. **h,** Normalized CC score distribution of sub-tomograms comparing with original and 36⁰ rotated sub-tomogram averages. **i,** Distribution of CC score differences for sub-tomograms comparing with original and 36⁰ rotated sub-tomogram averages.

**Extended Data Fig. 3.**
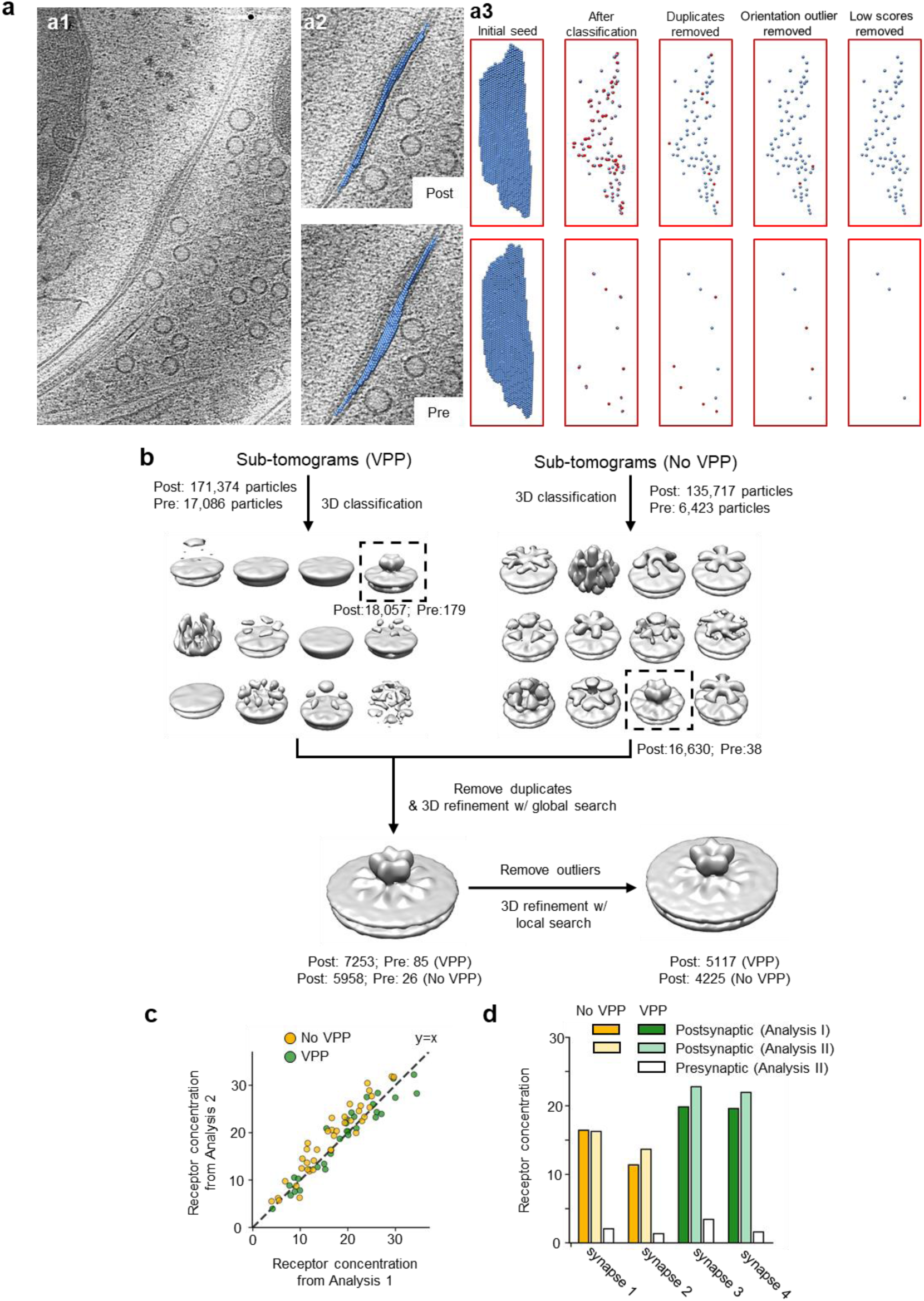
Identification of GABA_A_Rs using sub-tomograms mixed with sub-tomograms on presynaptic membrane. **a,** Steps for identifying GABA_A_Rs from sampling points mixed with sub-tomograms on presynaptic membrane. **a1,** Electron tomographic slice of an inhibitory synapse. **a2,** Electron tomographic slices superposed with sampling points on postsynaptic membrane (top) or presynaptic membrane (bottom). **a3,** Sub-tomogram sampling points on postsynaptic membrane (top) or presynaptic (bottom) after each step. Red points are sampling points that will be discarded in the next step. **b,** 3D classification and refinement of sub-tomograms on 72 postsynaptic membranes and 4 presynaptic membranes. **c,** Receptor concentration from Analysis II (analysis of sub-tomograms mixed with presynaptic sub-tomograms) verses receptor concentration from Analysis I (analysis of sub-tomograms without presynaptic sub-tomograms). **d,** Concentration of identified receptors on postsynaptic membranes in Analysis I, Analysis II, and falsely identified receptors on presynaptic membranes in Analysis 2 for the 4 selected synapses.

**Extended Data Fig. 4.**
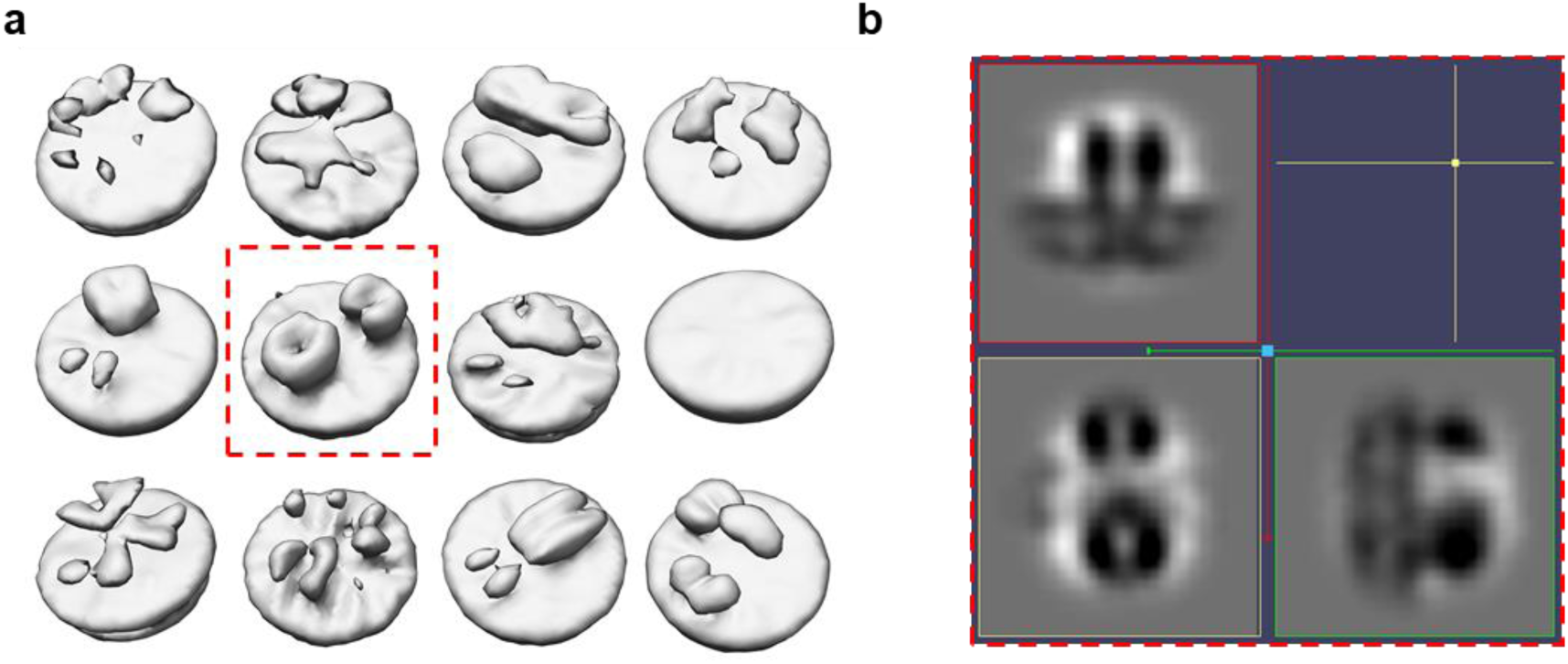
Classification of oversampled sub-tomograms without symmetry. **a,** Structures obtained from the 3D classification with VPP data. **b,** Orthogonal slice views of the structure boxed in (**a**).

**Extended Data Fig. 5.**
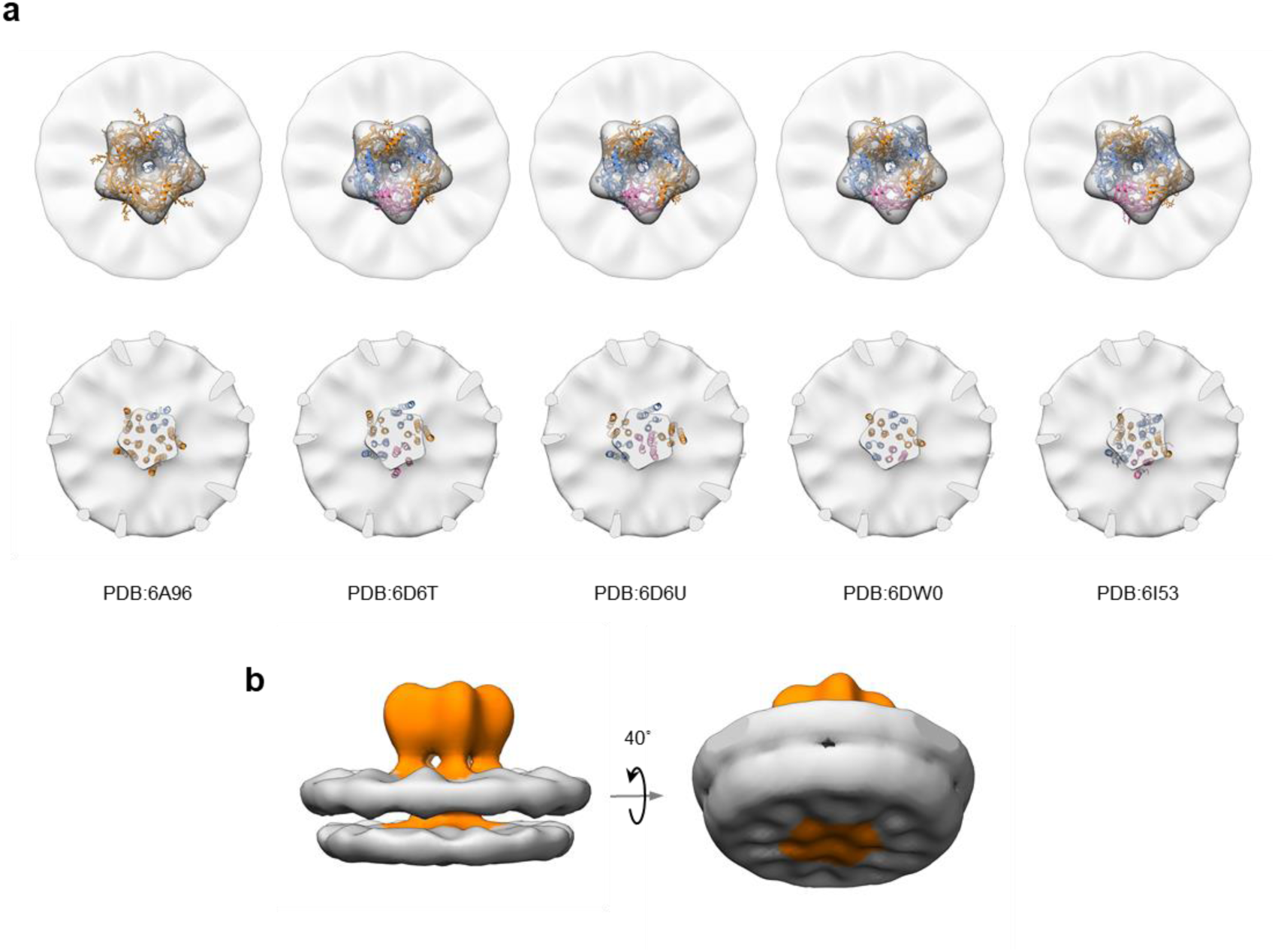
Structure features of sub-tomogram average of GABA_A_R. **a,** Density of the sub-tomogram average fitted with atomic models of different subunit compositions or conformations^26–29^. **b,** Left: Sub-tomogram average of GABA_A_R. Orange density is GABA_A_R density. Gray density is membrane bilayer. Right: Rotated view of sub-tomogram average of GABA_A_R displayed at low threshold.

**Extended Data Fig. 6.**
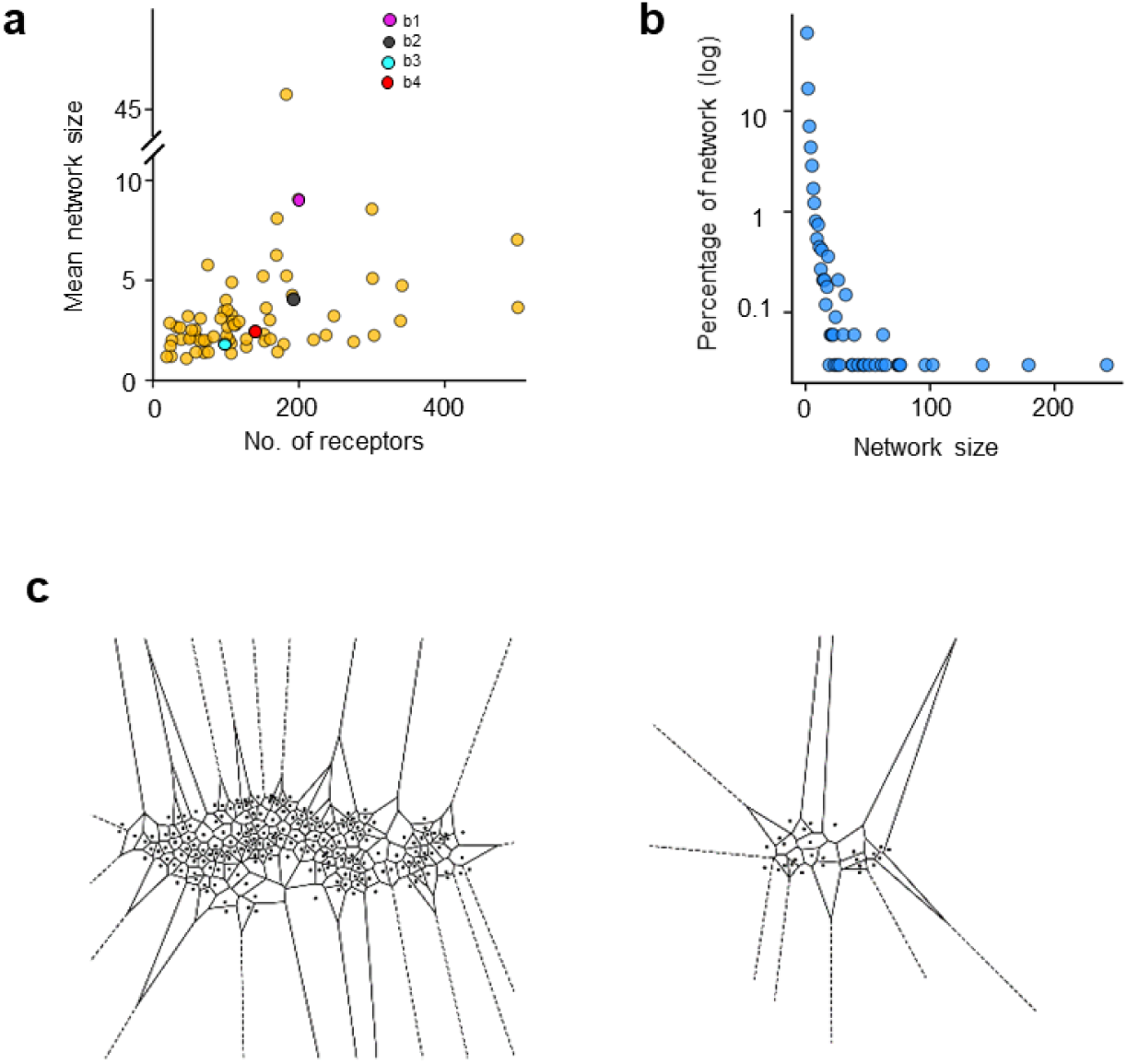
Two dimensional networks of GABA_A_Rs. **a,** Scattered plot of mean network size (number of receptors divided by number of networks) and number of receptors for each synapse. Colored dots (magenta, gray, cyan and red) correspond to the four synapses in Fig. 3b respectively. **b,** Distribution of network size. Y-axis is plot in logarithm scale. **c,** Examples of Voronoi tessellation of receptors on postsynaptic membrane. Black dots represent the localizations of GABA_A_Rs.

**Extended Data Fig. 7.**
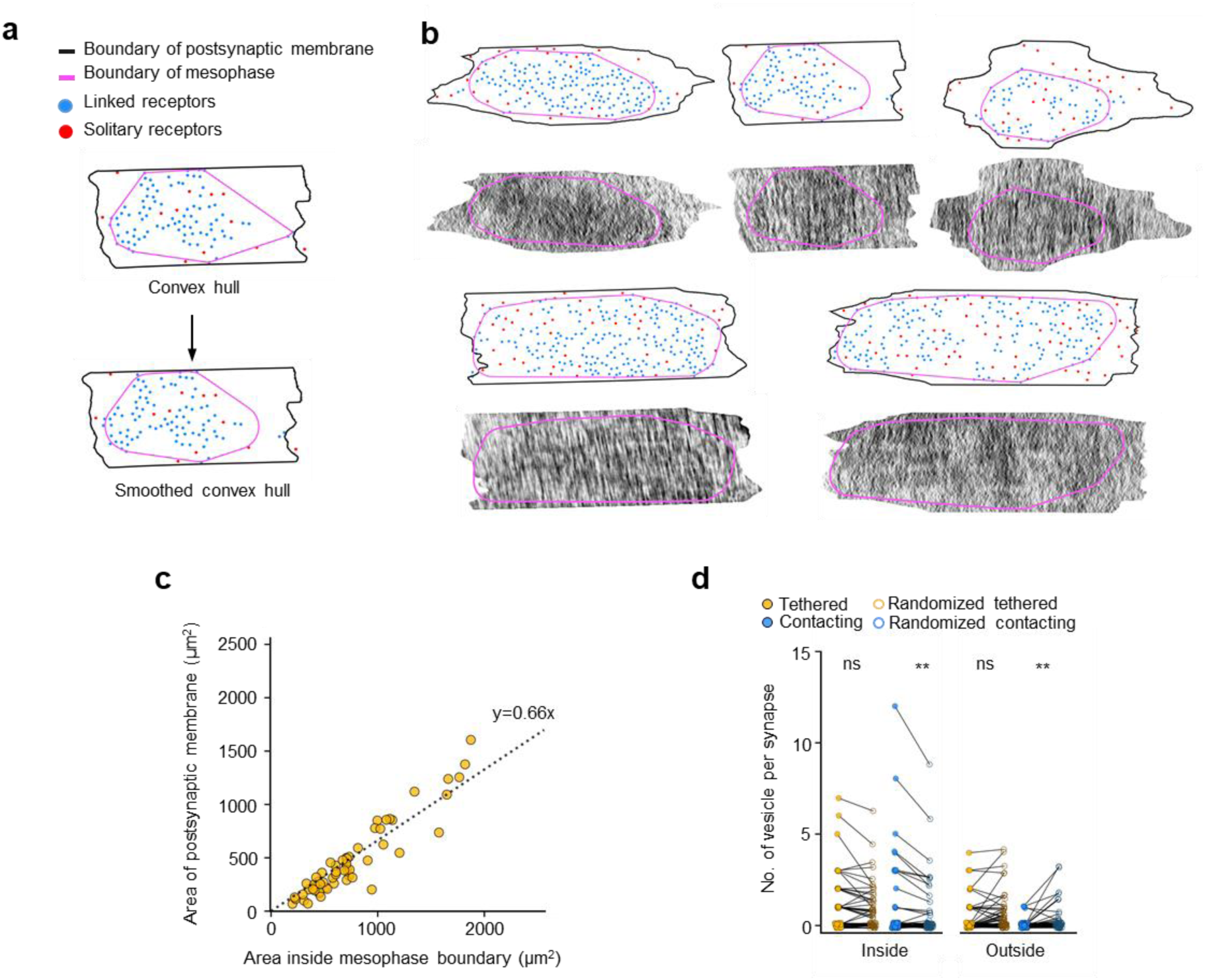
Mesophasic assembly of PSD. **a,** Example of convex hull and smooth convex hull of linked receptors. **b,** Examples of receptors distributions and 2D density profiles of scaffolding layer. **c,** Scatted plot of area inside mesophase boundary and area of postsynaptic membrane, fitted with a dashed line. **d,** Comparison between the number of either type of vesicles inside or outside of mesophase boundary with the corresponding number based on randomized vesicle distribution within the whole synapse (n=58 synapses). ns, P=0.20; **, P=0.005, paired two-tailed t-test.

**Extended Data Table 1.**
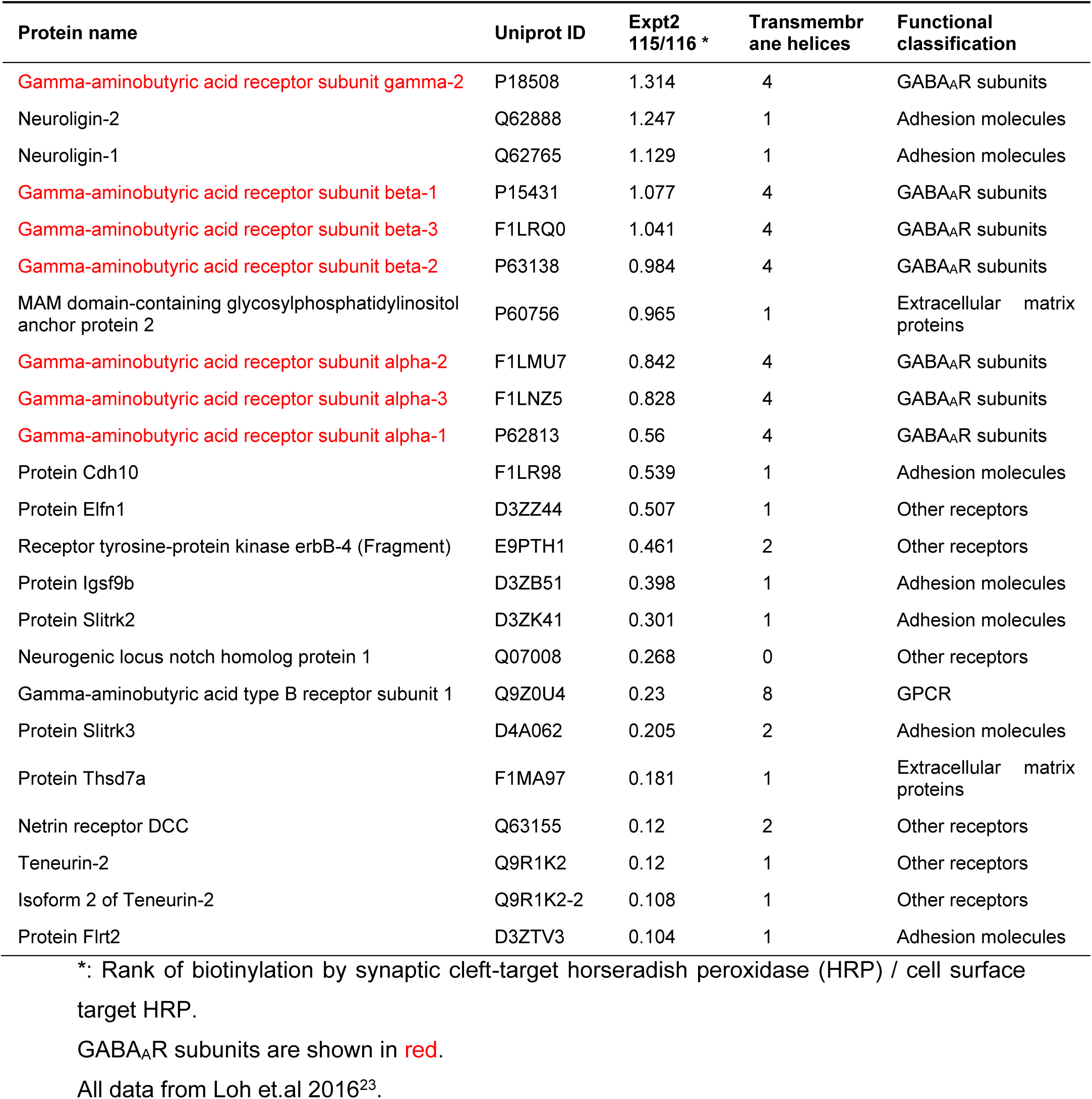
Proteomes on inhibitory postsynaptic membrane

## Supplementary

**Supplementary Video 1.**
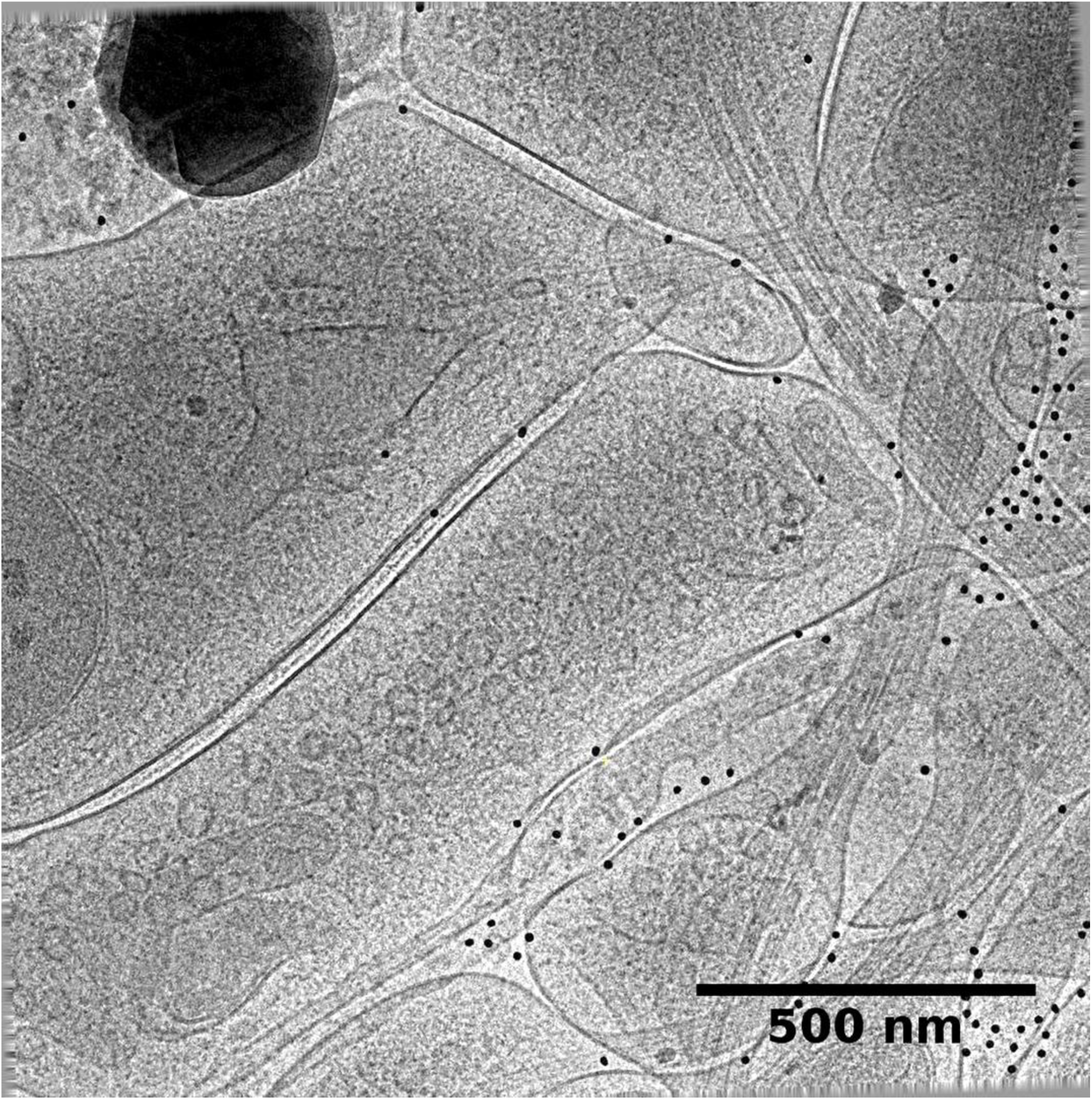
Tilt series of an inhibitory synapse. This video shows the tilt series of an inhibitory synapse (same data as in Fig. 1b) obtained using VPP, electron filter and counting technologies. Black dots of 15-nm diameter are gold beads used as fiducial markers for image alignment.

**Supplementary Video 2.**
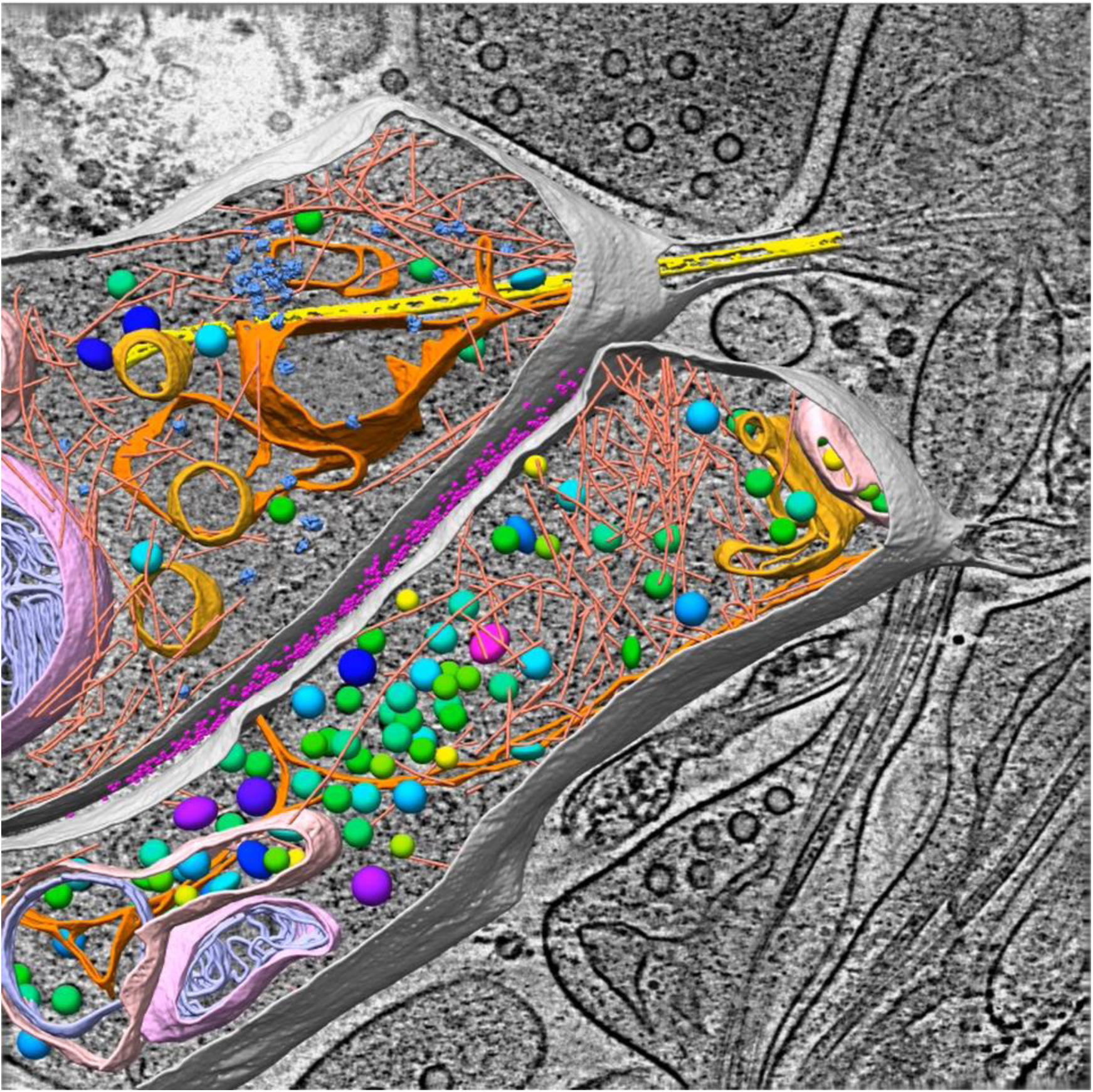
Structures of an inhibitory synapse. This video shows the tomogram of the same synapse as in Fig. 2b, displayed as z-stack and 3D surface rendering of the segmented structures. The structures were colored same as Fig. 1c and Fig. 1d.

## References

1 Eccles, J. C. The physiology of synapses. (Springers, 1964).

2 Sudhof, T. C. & Malenka, R. C. Understanding Synapses: Past, Present, and Future. Neuron 60, 469–476 (2008).

3 Mayford, M., Siegelbaum, S. A. & Kandel, E. R. Synapses and memory storage. Cold Spring Harbor perspectives in biology 4, doi:10.1101/cshperspect.a005751 (2012).

4 Sheng, M., Sabatini, B. L. & Sudhof, T. C. Synapses and Alzheimer’s disease. Cold Spring Harbor perspectives in biology 4, doi:10.1101/cshperspect.a005777 (2012).

5 Dosemeci, A., Weinberg, R. J., Reese, T. S. & Tao-Cheng, J. H. The Postsynaptic Density: There Is More than Meets the Eye. Frontiers in synaptic neuroscience 8, 23, doi:10.3389/fnsyn.2016.00023 (2016).

6 Liu, Y. T., Tao, C. L., Lau, P. M., Zhou, Z. H. & Bi, G. Q. Postsynaptic protein organization revealed by electron microscopy. Current opinion in structural biology 54, 152–160, doi:10.1016/j.sbi.2019.02.012 (2019).

7 Tao, C. L. et al. Differentiation and Characterization of Excitatory and Inhibitory Synapses by Cryo-electron Tomography and Correlative Microscopy. J Neurosci 38, 1493–1510, doi:10.1523/JNEUROSCI.1548-17.2017 (2018).

8 Valtschanoff, J. G. & Weinberg, R. J. Laminar organization of the NMDA receptor complex within the postsynaptic density. J Neurosci 21, 1211–1217 (2001).

9 Tang, A. H. et al. A trans-synaptic nanocolumn aligns neurotransmitter release to receptors. Nature 536, 210–214, doi:10.1038/nature19058 (2016).

10 Pennacchietti, F. et al. Nanoscale Molecular Reorganization of the Inhibitory Postsynaptic Density Is a Determinant of GABAergic Synaptic Potentiation. J Neurosci 37, 1747–1756, doi:10.1523/JNEUROSCI.0514-16.2016 (2017).

11 Mele, M., Leal, G. & Duarte, C. B. Role of GABAA R trafficking in the plasticity of inhibitory synapses. J Neurochem 139, 997–1018, doi:10.1111/jnc.13742 (2016).

12 Penn, A. C. et al. Hippocampal LTP and contextual learning require surface diffusion of AMPA receptors. Nature 549, 384–388, doi:10.1038/nature23658 (2017).

13 Chen, X. et al. Organization of the core structure of the postsynaptic density. Proc Natl Acad Sci U S A 105, 4453–4458, doi:10.1073/pnas.0800897105 (2008).

14 DeGiorgis, J. A., Galbraith, J. A., Dosemeci, A., Chen, X. & Reese, T. S. Distribution of the scaffolding proteins PSD-95, PSD-93, and SAP97 in isolated PSDs. Brain cell biology 35, 239–250, doi:10.1007/s11068-007-9017-0 (2006).

15 Sheng, M. & Kim, E. The postsynaptic organization of synapses. Cold Spring Harbor perspectives in biology 3, doi:10.1101/cshperspect.a005678 (2011).

16 Nair, D. et al. Super-resolution imaging reveals that AMPA receptors inside synapses are dynamically organized in nanodomains regulated by PSD95. J Neurosci 33, 13204–13224, doi:10.1523/JNEUROSCI.2381-12.2013 (2013).

17 MacGillavry, H. D., Song, Y., Raghavachari, S. & Blanpied, T. A. Nanoscale Scaffolding Domains within the Postsynaptic Density Concentrate Synaptic AMPA Receptors. Neuron 78, 615–622, doi:10.1016/j.neuron.2013.03.009 (2013).

18 Crosby, K. C. et al. Nanoscale Subsynaptic Domains Underlie the Organization of the Inhibitory Synapse. Cell reports 26, 3284-+, doi:10.1016/j.celrep.2019.02.070 (2019).

19 Zeng, M. et al. Reconstituted Postsynaptic Density as a Molecular Platform for Understanding Synapse Formation and Plasticity. Cell 174, 1172–1187 e1116, doi:10.1016/j.cell.2018.06.047 (2018).

20 Zeng, M. et al. Phase Transition in Postsynaptic Densities Underlies Formation of Synaptic Complexes and Synaptic Plasticity. Cell 166, 1163–1175 e1112, doi:10.1016/j.cell.2016.07.008 (2016).

21 Specht, C. G. et al. Quantitative nanoscopy of inhibitory synapses: counting gephyrin molecules and receptor binding sites. Neuron 79, 308–321, doi:10.1016/j.neuron.2013.05.013 (2013).

22 Miller, P. S. & Aricescu, A. R. Crystal structure of a human GABAA receptor. Nature 512, 270–275, doi:10.1038/nature13293 (2014).

23 Loh, K. H. et al. Proteomic Analysis of Unbounded Cellular Compartments: Synaptic Clefts. Cell 166, 1295–1307 e1221, doi:10.1016/j.cell.2016.07.041 (2016).

24 Nusser, Z., Hajos, N., Somogyi, P. & Mody, I. Increased number of synaptic GABA(A) receptors underlies potentiation at hippocampal inhibitory synapses. Nature 395, 172–177, doi:10.1038/25999 (1998).

25 Scheres, S. H. RELION: implementation of a Bayesian approach to cryo-EM structure determination. J Struct Biol 180, 519–530, doi:10.1016/j.jsb.2012.09.006 (2012).

26 Zhu, S. et al. Structure of a human synaptic GABAA receptor. Nature, doi:10.1038/s41586-018-0255-3 (2018).

27 Liu, S. et al. Cryo-EM structure of the human alpha5beta3 GABAA receptor. Cell Res 28, 958–961, doi:10.1038/s41422-018-0077-8 (2018).

28 Phulera, S. et al. Cryo-EM structure of the benzodiazepine-sensitive alpha1beta1gamma2S tri-heteromeric GABAA receptor in complex with GABA. eLife 7, doi:10.7554/eLife.39383 (2018).

29 Laverty, D. et al. Cryo-EM structure of the human alpha1beta3gamma2 GABAA receptor in a lipid bilayer. Nature 565, 516–520, doi:10.1038/s41586-018-0833-4 (2019).

30 Blanpied, T. A., Kerr, J. M. & Ehlers, M. D. Structural plasticity with preserved topology in the postsynaptic protein network. Proc Natl Acad Sci U S A 105, 12587–12592, doi:10.1073/pnas.0711669105 (2008).

31 Bak, P., Tang, C. & Wiesenfeld, K. Self-organized criticality: An explanation of the 1/fnoise. Physical review letters 59, 381–384, doi:10.1103/PhysRevLett.59.381 (1987).

32 Bormashenko, E. et al. Characterization of Self-Assembled 2D Patterns with Voronoi Entropy. Entropy-Switz 20, doi:10.3390/e20120956 (2018).

33 Limaye, A. V., Narhe, R. D., Dhote, A. M. & Ogale, S. B. Evidence for convective effects in breath figure formation on volatile fluid surfaces. Physical review letters 76, 3762–3765, doi:DOI 10.1103/PhysRevLett.76.3762 (1996).

34 Zuber, B. & Unwin, N. Structure and superorganization of acetylcholine receptor-rapsyn complexes. Proc Natl Acad Sci U S A 110, 10622–10627, doi:10.1073/pnas.1301277110 (2013).

35 Heuser, J. E. & Salpeter, S. R. Organization of acetylcholine receptors in quick-frozen, deep-etched, and rotary-replicated Torpedo postsynaptic membrane. The Journal of cell biology 82, 150–173, doi:10.1083/jcb.82.1.150 (1979).

36 Fernandez-Busnadiego, R. et al. Quantitative analysis of the native presynaptic cytomatrix by cryoelectron tomography. The Journal of cell biology 188, 145–156, doi:10.1083/jcb.200908082 (2010).

37 Zuber, B. & Lucic, V. Molecular architecture of the presynaptic terminal. Current opinion in structural biology 54, 129–138, doi:10.1016/j.sbi.2019.01.008 (2019).

38 Sola, M. et al. Structural basis of dynamic glycine receptor clustering by gephyrin. The EMBO journal 23, 2510–2519, doi:10.1038/sj.emboj.7600256 (2004).

39 Saiepour, L. et al. Complex role of collybistin and gephyrin in GABAA receptor clustering. J Biol Chem 285, 29623–29631, doi:10.1074/jbc.M110.121368 (2010).

40 Heine, M., Karpova, A. & Gundelfinger, E. D. Counting gephyrins, one at a time: a nanoscale view on the inhibitory postsynapse. Neuron 79, 213–216, doi:10.1016/j.neuron.2013.07.004 (2013).

41 Tretter, V. et al. Gephyrin, the enigmatic organizer at GABAergic synapses. Frontiers in cellular neuroscience 6, 23, doi:10.3389/fncel.2012.00023 (2012).

## References for Methods

42 Tao, C. L., Liu, Y. T., Zhou, Z. H., Lau, P. M. & Bi, G. Q. Accumulation of Dense Core Vesicles in Hippocampal Synapses Following Chronic Inactivity. Front Neuroanat 12, 48, doi:10.3389/fnana.2018.00048 (2018).

43 Sutton, M. A. et al. Miniature neurotransmission stabilizes synaptic function via tonic suppression of local dendritic protein synthesis. Cell 125, 785–799, doi:10.1016/j.cell.2006.03.040 (2006).

44 Sun, R. et al. An efficient protocol of cryo-correlative light and electron microscopy for the study of neuronal synapses. Biophysics Reports 5, 111–122, doi:10.1007/s41048-019-0092-4 (2019).

45 Kremer, J. R., Mastronarde, D. N. & McIntosh, J. R. Computer visualization of three-dimensional image data using IMOD. J Struct Biol 116, 71–76, doi:10.1006/jsbi.1996.0013 (1996).

46 Mastronarde, D. N. Automated electron microscope tomography using robust prediction of specimen movements. J Struct Biol 152, 36–51, doi:10.1016/j.jsb.2005.07.007 (2005).

47 Li, X. M. et al. Electron counting and beam-induced motion correction enable near-atomic-resolution single-particle cryo-EM. Nature methods 10, 584-+, doi:10.1038/nmeth.2472 (2013).

48 Rohou, A. & Grigorieff, N. CTFFIND4: Fast and accurate defocus estimation from electron micrographs. J Struct Biol 192, 216–221, doi:10.1016/j.jsb.2015.08.008 (2015).

49 Turonova, B., Schur, F. K. M., Wan, W. & Briggs, J. A. G. Efficient 3D-CTF correction for cryo-electron tomography using NovaCTF improves subtomogram averaging resolution to 3.4A. J Struct Biol 199, 187–195, doi:10.1016/j.jsb.2017.07.007 (2017).

50 Pettersen, E. F. et al. UCSF Chimera--a visualization system for exploratory research and analysis. J Comput Chem 25, 1605–1612, doi:10.1002/jcc.20084 (2004).

51 Hrabe, T. et al. PyTom: a python-based toolbox for localization of macromolecules in cryo-electron tomograms and subtomogram analysis. J Struct Biol 178, 177–188, doi:10.1016/j.jsb.2011.12.003 (2012).

52 Mattei, S., Glass, B., Hagen, W. J. H., Krausslich, H. G. & Briggs, J. A. G. The structure and flexibility of conical HIV-1 capsids determined within intact virions. Science 354, 1434–1437, doi:10.1126/science.aah4972 (2016).

53 Navarro, P. P., Stahlberg, H. & Castano-Diez, D. Protocols for Subtomogram Averaging of Membrane Proteins in the Dynamo Software Package. Front Mol Biosci 5, 82, doi:10.3389/fmolb.2018.00082 (2018).

54 Zivanov, J. et al. New tools for automated high-resolution cryo-EM structure determination in RELION-3. eLife 7, doi:10.7554/eLife.42166 (2018).

55 Rosenthal, P. B. & Henderson, R. Optimal determination of particle orientation, absolute hand, and contrast loss in single-particle electron cryomicroscopy. J Mol Biol 333, 721–745 (2003).

